# Translational antagonism enables bet-hedging in yeast via mitochondrial bistability

**DOI:** 10.64898/2026.03.30.715437

**Authors:** Piyush Nanda, Andrew W. Murray

**Affiliations:** Program in Biological and Biomedical Sciences, Harvard Medical School, Boston, MA 02115, USA; Department of Molecular and Cellular Biology, Harvard University, Cambridge, MA 02138, USA

## Abstract

We investigated the molecular basis of bet-hedging in budding yeast, *Saccharomyces cerevisiae*. Fast glycolytic growth generates two cell states: fermenting arrestors and respiring recoverers, of which only recoverers resume growth when shifted to a respiratory carbon source. Single-cell metabolic biosensors show that a bistable switch in mitochondrial activity produces these states: mitochondrial membrane potential drives import of positively charged, nuclear-encoded proteins of the mitochondrial ribosome, and mitochondrial ribosomes synthesize electron transport chain subunits that sustain the potential, completing a positive feedback loop. The ratio of mitochondrial to cytoplasmic protein synthesis governs switch dynamics: the effect of slowing mitochondrial translation is rescued by inhibiting cytosolic translation. Reducing mitochondrial translation reconstitutes bistability in evolutionarily distant fission yeast, suggesting a conserved mechanism for diversifying single-cell ATP production strategies.

## Introduction

In the presence of oxygen and glucose, cells face a fundamental decision: whether to ferment or respire glucose to produce ATP. Fast growth on abundant glucose leads to increased fermentation^1^ and reduced respiration despite the presence of oxygen in bacteria^2,3^, yeast^3^, and cancer cells^4^. As glucose availability decreases, however, respiration becomes essential to meet the energetic demands of growth^1,5^. In yeast, during fermentative growth, an estimated ∼10^8^ glucose molecules enter the cell every second, sustaining an ATP pool of ∼10^7^ molecules that is rapidly turned over to power all growth-associated processes. Thus, each cell generates and uses its entire ATP pool roughly every 50-100 milliseconds, making cellular ATP levels vulnerable to environmental glucose fluctuations. This rapid turnover raises the question of how individual cells buffer changes in the availability of ATP as they switch between respiration and fermentation in unpredictable environments.

In *Saccharomyces cerevisiae*^6^, abrupt glucose deprivation exposes two pre-existing cell states: arrestors and recoverers. Recoverers divide slower than arrestors during fermentative growth on high glucose but can transition to alternative carbon sources whereas arrestors cannot. During steady-state growth on glucose, yeast cells spontaneously switch between the two states to maintain roughly 25% recoverers on high glucose. Slow switching, relative to growth, creates lineages that share the same metabolic fate.

To uncover the mechanism that produces the two states on glucose, we imaged the metabolic states of individual cells in a microfluidic system during the transition from glucose to galactose or ethanol, substrates that cells respire rather than fermenting. By combining single-cell metabolic measurements with physiological perturbations, we uncovered a bistable switch in mitochondrial protein synthesis that generates the recoverer and arrestor states during growth on glucose. The steady-state recoverer fraction increases as cells are grown in lower glucose concentrations and this response is modulated through sensors that regulate the pathway by which Ras activity regulates the activity of cyclic AMP-dependent protein kinase A (cAMP/PKA). Bistability arises from a non-linear positive feedback loop that reflects competition between mitochondrial and cytoplasmic protein synthesis: 1) recoverers use the large potential difference across the mitochondrial membrane to drive the import of the protein components of the mitochondrial ribosome, which are translated in the cytoplasm, and 2) the mitochondrial ribosomes translate the components of the electron transport chain that are encoded in the mitochondrial DNA (mtDNA), and 3) electron transport sustains the membrane potential by pumping protons across the inner membrane. In arrestors, the lower potential difference decreases mitochondrial protein synthesis leading to less electron transport and maintaining the lower potential. Fast growth dilutes the mitochondrial proteome and consequently biases switching towards arrestors. Tuning mitochondrial translation in *Schizosaccharomyces pombe*, which normally only shows the recoverer phenotype, reconstitutes bistability, suggesting a conserved mechanism for switching between fermentation and respiration.

### External glucose levels tune the recoverer fraction in the population

We followed individual cells to determine their response to switching from steady state growth in different glucose concentrations to 1% (v/v) ethanol or 2 % (w/v) galactose, both non-fermentable carbon sources. We used a fully automated analysis pipeline for inferring the recoverer fraction from time-lapse movies (**Figure S1**). The steady state recoverer fraction increases as the pre-shift glucose concentration decreases (**Figure 1A**). This environmental response depends on the activation of adenylyl cyclase by Ras leading to increased cyclic AMP and cAMP-dependent protein kinase activity^7^. Hyperactivation of Ras^7^, by removing its inactivators (Ira1 and Ira2), eliminates recoverers from the population for all glucose concentrations (**Figure 1A**). Conversely, reducing the activity of cAMP-dependent protein kinase increases the recoverer fraction in a dose-dependent manner (**Figure S2**). Changing the post-shift carbon source between galactose and ethanol, metabolized by different pathways, does not significantly alter the fraction of recoverers (**Figure S3**), arguing that it is the state of the cell before the environmental perturbation, not the new environment, that determines its response to the disappearance of glucose. Together, our results suggest that a mechanism generates spontaneous slow switching and sets the metabolic state of arrestors and recoverers during fermentative growth before the shift. To understand this mechanism, we measured the energetic states of the two subpopulations as they switch from glucose to galactose using PercevalHR^8^, a sensor for intracellular ATP:ADP ratio.

**Figure 1.**
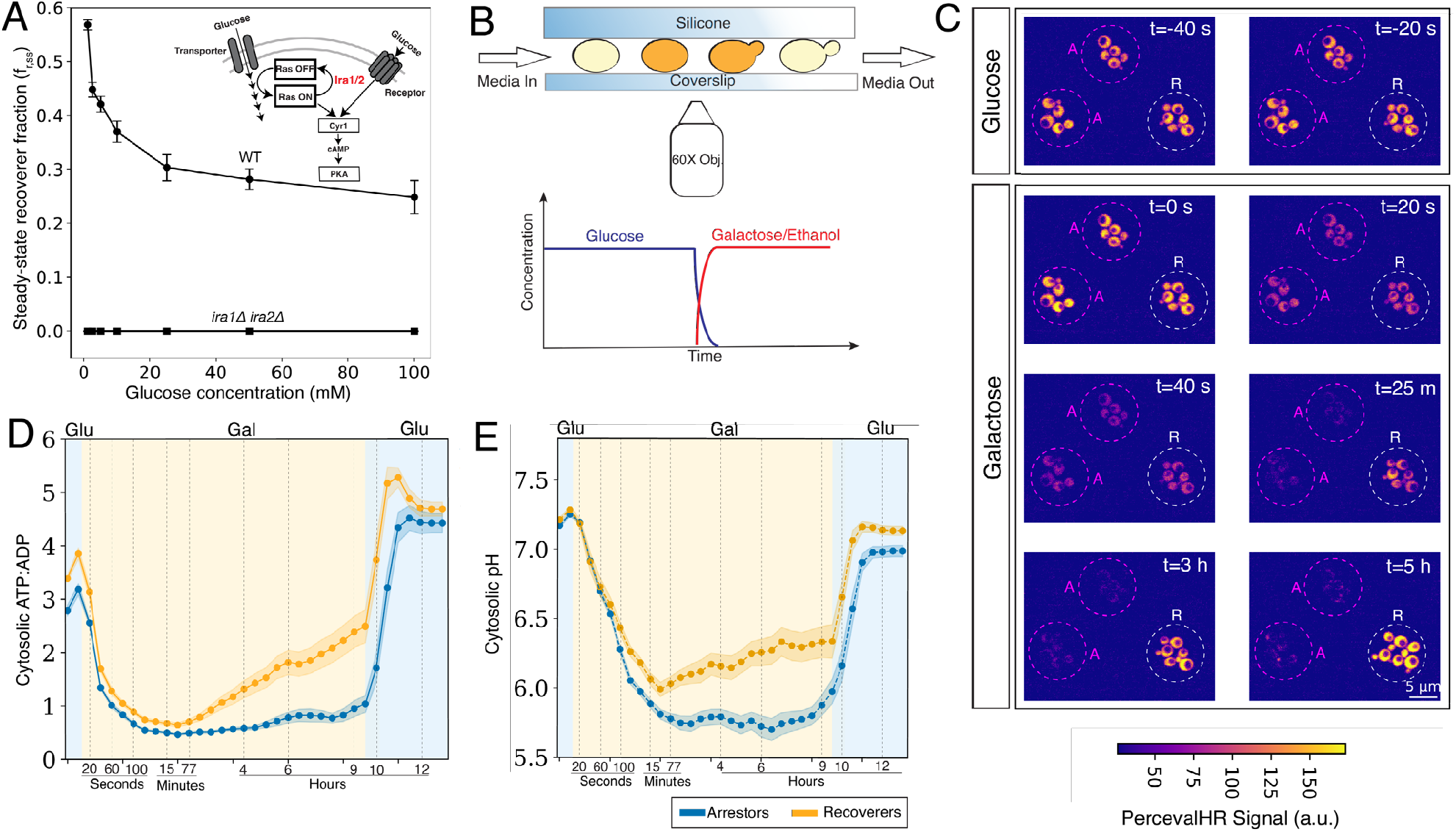
Differences in the instantaneous response to glucose deprivation define the two cell states (A) The steady-state fraction of recoverers is set by the preshift glucose concentration and is under genetic control. Ira1 and Ira2 are paralogous Ras GTPase activating proteins. (B) A microfluidic set-up for using fluorescent biosensors to measure cytosolic ATP:ADP ratio and pH following glucose deprivation. (C) Changes in cytosolic ATP:ADP ratio, measured using a calibrated PercevalHR sensor, following glucose deprivation. Three microcolonies (circled) are shown: two are arrestors (A) and the third composed of recoverers (R).(D) Extended measurement of cytosolic ATP:ADP in arrestors (n=31) and recoverers (n=24) on various timescales (seconds, minutes, and hours) in a single experiment. The blue wash indicates when cells were exposed to glucose, the yellow when they were exposed to 2% galactose. (E) Measurement of cytosolic pH in arrestors (n=35) and recoverers (n=35) using super-ecliptic pHluorin.

### Rapid metabolic response to glucose removal sets recoverers on a growth-recovery trajectory

We asked if the difference between arrestors and recoverers’ instantaneous response to glucose deprivation sets their fates. We calibrated (Methods) PercevalHR to continuously measure absolute ATP: ADP ratios (**Figure S4A-D**) in cells as their environment switched from glucose to galactose in a microfluidic set-up (**Figure 1B**). By adjusting the frequency of imaging in real-time, we captured the ATP:ADP ratio dynamics from seconds to hours in individual cells. The ATP:ADP ratio fell rapidly in arrestors and did not recover despite prolonged incubation in the presence of galactose. Even in the first few seconds after the shift, ATP:ADP falls less in recoverers, and they then slowly reach ATP:ADP ratios that allow them to resume growth in a few hours (**Figure 1C-D**). We simultaneously imaged the cytosolic pH using a strain expressing super-ecliptic pHluorin (SEP) along with the PercevalHR expressing cells in the same microfluidic chip (**Figure S4A**). Cytosolic pH recovers on similar timescales as the ATP:ADP ratios in recoverers but not in arrestors (**Figure 1E**). On refeeding glucose, arrestors rapidly resume their metabolism and start proliferating, showing they remain viable despite low ATP:ADP ratios and cytosolic pH (**Figure 1D-E**).

The initial ATP:ADP dynamics following the shift are well described by a simple model (**Figure S4E**) in which recoverers continue to produce ATP at a reduced rate within a few seconds after the shift. The initial, nearly exponential ATP:ADP decay rate, determined by ATP consumption exceeding production, is similar in both states (*t_half,ATP:ADP_* = 0.19 min *or* 11 *secs*), but recoverers maintain a higher asymptotic ATP:ADP at the lowest point in the trajectory (1.03 in recoverers vs 0.81 in arrestors). We hypothesized that recoverers, but not arrestors, were primarily respiring on high glucose, which allows them to continue producing ATP in the absence of glucose, using intracellular reserves or slowly consuming the post-shift carbon source—both of which require respiration. To test this, we measured two aspects of metabolism in single cells before the transition to galactose (**Figure 2A**): i) glycolytic flux by using a sensor of Fructose 1,6 bisphosphate (FBP), that exploits a transcriptional repressor that is allosterically modulated by FBP and whose response is proportional to glycolytic flux^9^ ii) membrane potential across mitochondrial inner membrane (ΔΨ) as reported by the import of Tetramethyl rhodamine methyl ester (TMRM^10^).

**Figure 2.**
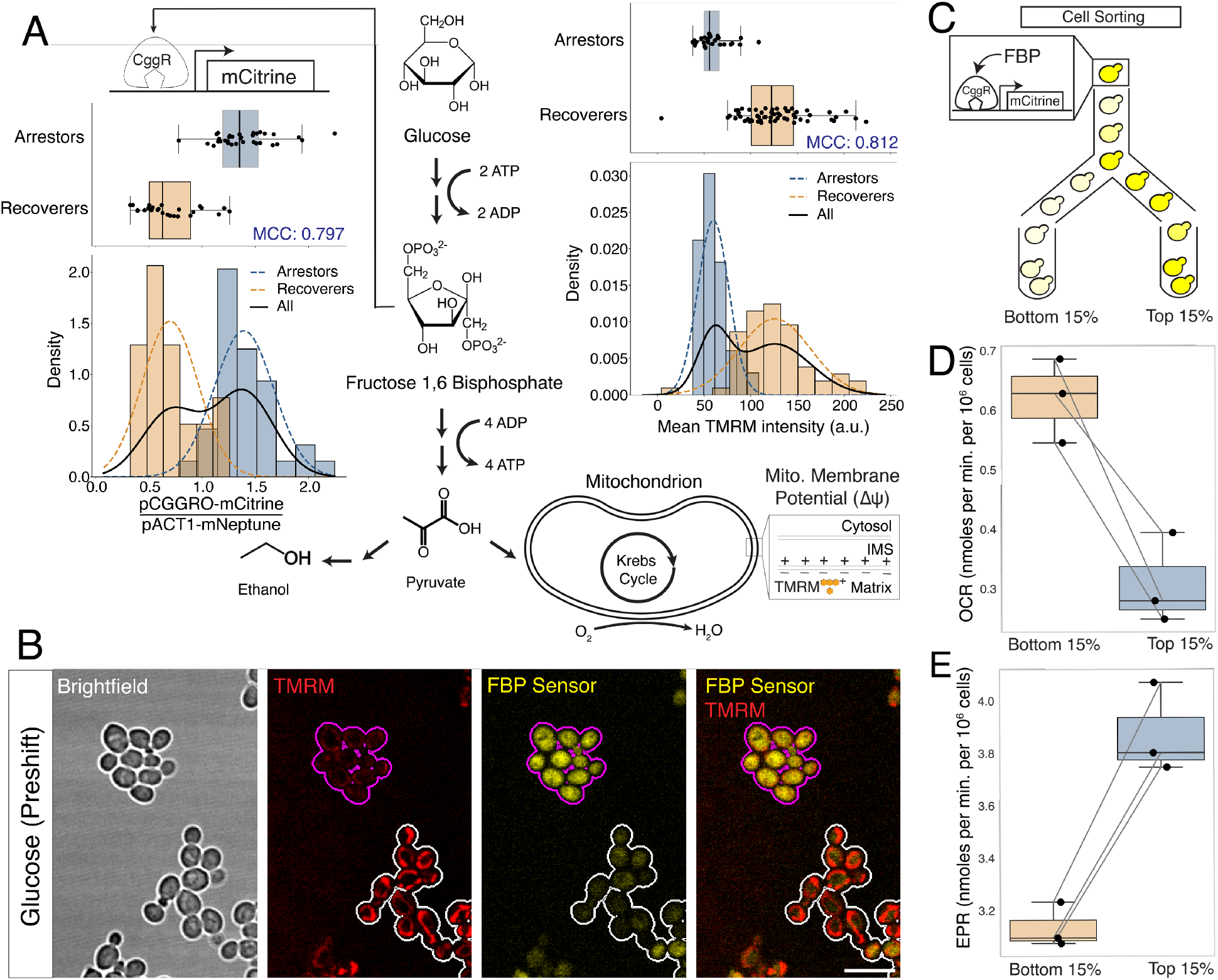
Arrestors ferment and recoverers respire prior to glucose removal (A) Sensors for fructose 1,6-bisphosphate (FBP) and mitochondrial membrane potential measurement distinguish respiration and fermentation. The center panel shows the key steps in the fermentation or respiration of glucose. The left panel shows the level of mCitrine (normalized using mNeptune driven from the constitutive *ACT1* promoter) expressed from a promoter (pCGGRO) that is repressed by an FBP-binding protein (CggR) in two ways: the upper panel shows the values of individual cells while the lower panel shows histograms and distributions. The right panel shows the mitochondrial level of a positively charged small molecule (tetramethyl rhodamine, TMRM) in arrestors and recoverers, with the data for individual cells and histograms of the distributions (bottom panel). (B) The signal from the FBP sensor (yellow) and TMRM accumulation (red) in arrestors and recoverers. The cells were assigned their states based on their recovery phenotype after shifting them to 2% galactose. White boundaries: recoverers; Magenta boundaries: arrestors. (C) Cartoon of FACS sorting of the top 15% and bottom 15% cells expressing the FBP sensor in 0.05% glucose to generate arrestor and recoverer enriched sub-populations to measure the rates of oxygen consumption and ethanol production. Measurement of oxygen consumption (D) and ethanol production rates (E) for the two sorted fractions (n=3 independent experiments; the solid lines connect the measurements from a single experiment).

### Pre-starvation respiro-fermentative metabolism predicts the single-cell metabolic fate

We asked if recoverers show decreased FBP levels and increased mitochondrial membrane potential (ΔΨ) relative to arrestors, which would confirm their heightened respiratory activity before glucose deprivation. We quantified FBP levels, proxying glycolytic flux, by normalizing the mCitrine level controlled by the FBP sensor with constitutively expressed mNeptune fluorescent protein (**Figure 2A**). Consistent with our hypothesis, recoverers had lower FBP levels than arrestors, suggesting lower glycolytic flux, and a higher mitochondrial membrane potential, suggesting a higher respiratory activity. The metabolic states are stably maintained over several cell divisions (**Figure 2B**) consistent with the slow stochastic switching between the two states, we observed previously^6^. By scoring whether cells recovered after transition to galactose, we determined how well the FBP and TMRM signals predicted the fate of the two states. We used the Matthews Correlation Coefficient (MCC) to assess the predictive power of the FBP level and TMRM on glucose in determining the arrestor vs recoverer fate after transition to galactose. An MCC value of 1 indicates that the pre-shift reporter signal perfectly predicts the fate, whereas a value of 0 indicates no predictive power. The FBP signal is bimodal during growth on glucose and correctly predicts the fate of the individual cells on shift to galactose (MCC=0.797; **Figure 2A and S5A-B**). A strain lacking the FBP-responsive transcriptional repressor, thus decoupling the production of mCitrine from the FBP levels, showed no difference between arrestors and recoverers (**Figure S5C**). The TMRM signal also accurately predicts the fate of the two states (MCC=0.812; **Figure 2A and S6A-C**). These observations are consistent with a model in which arrestors and recoverers divert different fractions of the glycolytic flux into respiration and fermentation despite existing in the same, well-mixed environment before the shift.

FBP and TMRM measurements suggest different extents of fermentation and respiration in arrestors and recoverers but don’t directly measure either activity. We used Fluorescence Activated Cell Sorting (FACS) on the signal from the FBP sensor to sort arrestors (top 15%) and recoverers (bottom 15%) from a population growing on 0.05% glucose (**Figure 2C**). We then measured oxygen consumption (respiration) and ethanol production (fermentation) in these populations while they were still growing on glucose. Recoverers had higher oxygen consumption (**Figure 2D**) and lower ethanol production relative to the arrestors (**Figure 2E**). This observation demonstrates that arrestors and recoverers rely differentially on fermentation and respiration arguing that some circuit, whose state can be propagated for multiple cell divisions, establishes the cellular energy generation mode in the two states.

What molecular circuitry could generate the epigenetic inheritance of the arrestor and recoverer metabolic states? One simple mechanism is a positive feedback loop that creates a bistable switch which locks the cells in either of the two metabolic modes. We used our observations and the literature to propose a feedback loop that has two essential components: 1) increasing the magnitude of the mitochondrial membrane potential increases mitochondrial protein synthesis and 2) increasing mitochondrial protein synthesis increases the electron transport that produces the mitochondrial membrane potential. Mitochondrial membrane potential is responsible for import of proteins into the matrix through the TIM/TOM translocase complex^11,12^ as well as ATP synthesis. The proteins of the mitochondrial ribosome are encoded in the nucleus, translated in the cytoplasm, and must be imported into mitochondria where they assemble on the mitochondrially encoded and transcribed ribosomal RNA. Once assembled, mitochondrial ribosomes translate the mitochondrially encoded components of the electron transport chain and ATP synthase including three subunits of cytochrome oxidase (complex IV), the last step of electron transport. Mitochondrial ribosomal proteins are among the most highly charged polypeptides imported into mitochondria (**Figure S7A**) suggesting that their import should be strongly dependent on the charge across the inner mitochondrial membrane. To support this hypothesis, we used differently charged variants of GFP (supercharged GFP) to show that polypeptide charge density directly affects mitochondrial protein import rates (**Figure S7B-D**). Because the mitochondrial ribosomal proteins are positively charged, their import should be accelerated by the mitochondrial membrane potential due to an electrophoretic drive i.e. movement of charged molecules in an electric field. Because mitochondrial translation produces essential subunits of Complex IV and the activity of this complex sustains the mitochondrial membrane potential, it produces simple positive feedback (**Figure 3A**) that can create^13^ bistability. Low mitochondrial translation activity will lead to reduced membrane potential which leads to failure in ribosomal protein import, trapping cells in the arrestor state. Conversely, high mitochondrial translation will maintain the membrane potential sustaining the ribosomal protein import and propagating the recoverer state. Because of the self-reinforcing nature of the positive feedback the metabolic state will be maintained despite cell division. Occasional stochastic switching due to copy number fluctuations in components of the circuit would allow cells to switch between the two states.

**Figure 3.**
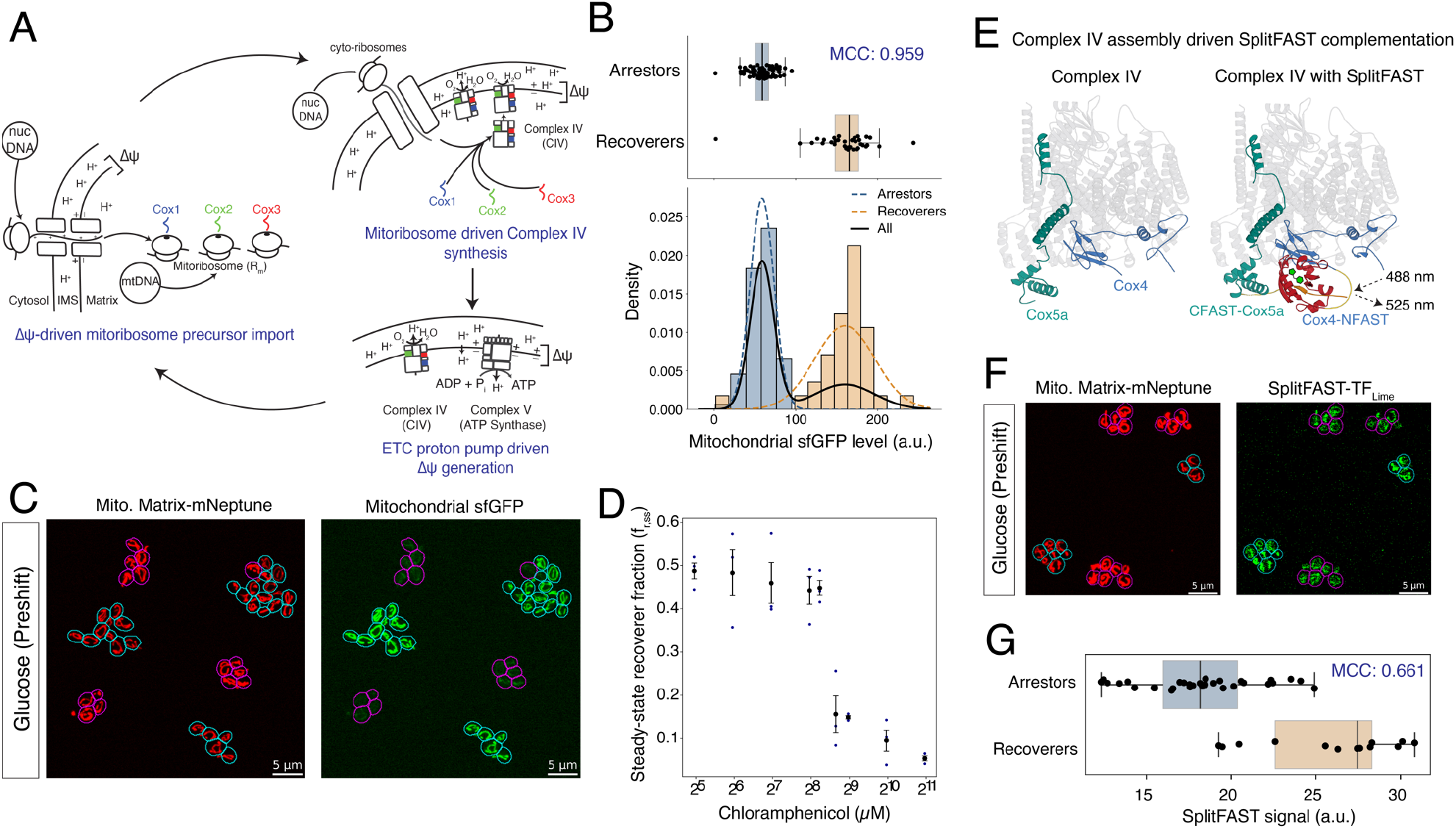
Mutual positive feedback between mitochondrial protein synthesis and membrane potential generates differential mitochondrial translation. (A) Membrane potential driven import of nuclear-encoded mitoribosome subunits and mitoribosome driven maintenance of membrane-potential generates a positive feedback loop. (B) Distribution of fluorescence intensity from a mtDNA-encoded superfolder GFP in arrestors and recoverers. The mitochondria were segmented using an independent marker (mNeptune) constitutively localized to the mitochondrial matrix using the PreSu9 pre-sequence. Arrestor and recoverer states were assigned based on their recovery after shift to 2% galactose. The upper panel shows the values of individual cells the lower, the distributions. (C) Representative image of cells expressing mtDNA encoded GFP (Green) based on their mitochondrial translation activity. A constitutively cytoplasmically translated mNeptune was targeted to the matrix using the PreSu9 pre-sequence. White boundaries: recoverers; Magenta boundaries: arrestors. (D) Changes in the steady state recoverer fraction in response to the indicated dose of the mitochondrial translation elongation inhibitor chloramphenicol (note the logarithmic scale) during growth on 0.05% glucose. (E) A reporter for assembly of complex IV based on fluorogen (HMBR/TF_Lime_) activation following SplitFAST reconstitution. (F) and (G) Split FAST-fluorogen signal in arrestors and recoverers growing on 0.2% glucose. The mitochondria were segmented using an independent marker (mNeptune) constitutively localized to mitochondrial matrix. Arrestor and recoverer states were assigned based on their recovery after shift to 2% galactose.

### Membrane-potential driven import of mitochondrial ribosomal proteins creates a positive feedback

To test whether arrestors and recoverers differ in mitochondrial translation, we used a yeast strain with a superfolder GFP gene integrated in the mitochondrial DNA^14^ (sfGFP^mt^). Because the sfGFP^mt^ mRNA is in the mitochondria matrix and can only be translated by mitochondrial ribosomes, the fluorescence signal reports mitochondrial translation activity. We found a bimodal distribution of fluorescence in the mitochondria which accurately predicted (MCC=0.959) whether cells would score as arrestors or recoverers (**Figure 3B-C**). Arrestors show a very low level of fluorescence despite possessing normal tubular mitochondrial structure as reported by a cytoplasmically synthesized mNeptune targeted to the mitochondrial matrix. Chloramphenicol selectively targets mitochondrial translation elongation without interfering with cytosolic translation^15^ suggesting that it should bias the switch in favor of arrestors. Titrating chloramphenicol produced a dose-dependent reduction in the steady state recoverer fraction, suggesting the mitochondrial translation rate is a control parameter of the circuit producing the two metabolic states (**Figure 3D**).

Our model predicts that differences in mitochondrial translation rate are reflected in the differences in complex IV assembly, which sustains the membrane potential. To measure assembled Complex IV, we used the Split-FAST^16^ system, which produces a fluorescent signal only when its two components are close enough to bind to each other. Based on the yeast complex IV structure^17^, we tagged the nuclear-encoded subunits Cox4 and Cox5a with the N and C terminal fragments of the FAST protein (**Figure 3E**) respectively. Stable assembly of Complex IV brings the two fragments close together reconstituting FAST (**Figure S8A**), which binds and increases the fluorescent yield of an exogenously supplied small molecule, 4-hydroxy-3-methylbenzylidene rhodanine (HMBR)/TF_lime_. Removal of Complex IV subunits or chaperones involved in its assembly significantly reduces the signal (**Figure S8B**) and the uptake of the dye is not reduced upon depolarization using a protonophore (**Figure S8C**). As our model predicts, the signal for Complex IV assembly is stronger in recoverers than in arrestors (MCC=0.661; **Figure 3F-G**). This difference is consistent with the difference in oxygen consumption, the catalytic reaction performed by Complex IV, between arrestors and recoverers. Our results establish that single yeast cells, in the same environment, adopt alternative metabolic states due to positive feedback between the mitochondrial translation that produces Complex IV and the membrane potential generated by Complex IV and the rest of the electron transport chain.

Hysteresis, or history dependence, implies bistability^18^. Because positive feedback reinforces each state, the threshold stimulus needed to switch between states depends on the system’s current state. To test for hysteresis, we exploited the ability of chloramphenicol to reduce the recoverer fraction by inhibiting mitochondrial protein synthesis: cells that start as arrestors (high chloramphenicol) should show different responses to intermediate chloramphenicol concentrations than those that start as recoverers (no chloramphenicol) (**Figure 4A**). We moved these two populations of cells to media containing intermediate concentrations of chloramphenicol, and continued culturing them for up to 18 hours before measuring the recoverer fraction. The dose-response to chloramphenicol depended on whether the populations began as arrestors or recoverers: for example, at 512 µM chloramphenicol, recoverers remained recoverers and arrestors remained arrestors (**Figure 4A**). Because the cells were dividing during the experiment and the effect continues for up to 18 hours, it originates from the slow relaxation of the population due to the underlying bistability that maintains cells as arrestors or recoverers.

**Figure 4.**
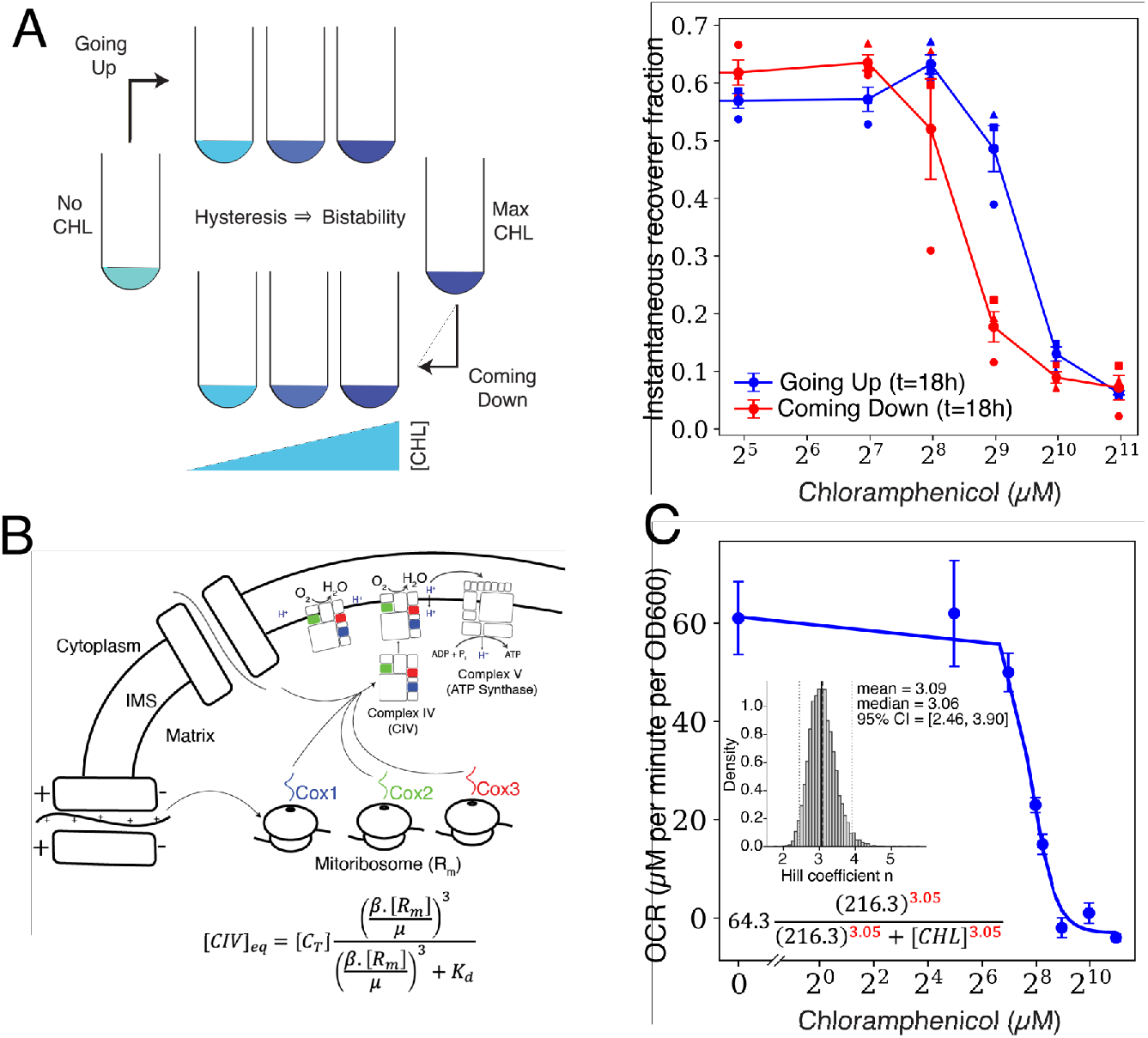
Ultrasensitive assembly of complex IV driven by the 3 mitochondrially encoded and translated subunits converts positive feedback into bistability. (A) Hysteresis in the fraction of recoverers on chloramphenicol titration. Cells grown in absence of chloramphenicol (going up) and 2 mM chloramphenicol (coming down) on 0.05% glucose were shifted to intermediate concentrations of chloramphenicol for 18 hours (Left). Experimental results from the hysteresis experiments on 0.05% glucose (Right). (B) A biophysical model for the assembly of complex IV co-operatively by mitoribosome synthesized subunits and nuclear-encoded subunits. (C) Measurement of oxygen consumption rate after chloramphenicol titration on 0.05% glucose. The oxygen uptake rate was measured using a modified oxoplate assay (methods). A Hill function was fit to the data to estimate the Hill exponent (blue).

### Ultrasensitive assembly of cytochrome oxidase converts positive feedback into a bistable switch

Positive feedback cannot produce bistability without non-linear interactions between components of the circuit^19^. Bimodality in mitochondrial membrane potential and translation and the hysteretic response to chloramphenicol all suggest unidentified non-linearities. We examined the details of the positive feedback circuit for features that could inject non-linearity into the circuit that connects mitochondrial translation to the generation of membrane potential. We noticed that Complex IV contains 3 mitochondrially (Cox1, Cox2 and Cox3) and 9 nuclear-encoded subunits^20^. If production of the three mitochondrially encoded subunits controls the rate of Complex IV assembly, its production will be ultra-sensitive to the rate of mitochondrial protein synthesis with a Hill exponent of 3 (**Figure 4B**), the number of mitochondrially encoded subunits.

We tested our quantitative prediction about the response to chloramphenicol by measuring the catalytic activity of Complex IV through the oxygen consumption rate: we found the Hill coefficient computed from the data (*n*=3.05±0.4), is statistically indistinguishable from the predicted number, *n=3* (**Figure 4C**). The split-FAST signal for the Complex IV assembly also produced a Hill exponent (*n*=3.45±0.28) close to the prediction (**Figure S9A-B**). Because it is hard to experimentally alter the predicted Hill coefficient, we cannot formally exclude the possibility that the agreement between predicted and measured values is coincidental. Nevertheless, the close agreement supports a simple biophysical model based on the contribution of mitochondrially synthesized components to the cooperative assembly of Complex IV.

Based on our experiments, we produced a simple biophysical model of the bistable circuit that connects potential-driven protein import, mitochondrial translation, and electron transport. By non-dimensionalization, we could reduce the predicted fraction of recoverers to an equation that contains a single dimensionless parameter that absorbs all the rate constants, charges and the concentrations (**Figure 5A**). This feedback strength parameter (*p*) and the Hill exponent, *n*=3, set the regime where a population contains both arrestors and recoverers. A higher parameter value biases switching towards recoverers. The rate constant of mitochondrial translation appears in the parameter’s numerator, and the denominator contains the specific growth rate of the cell, which is set by the cytosolic translation rate. Intuitively, these two forms of translation compete. If cells are growing fast, they will need more mitochondrial ribosomes to make the components needed to maintain electron transport and remain as recoverers: slowing mitochondrial protein synthesis will make it easier for recoverers to switch to arrestors and slowing cytoplasmic protein synthesis will have the opposite effect. Thus, slowing down cell growth by inhibiting cytoplasmic ribosomes should overcome the ability of chloramphenicol to reduce the recoverer fraction (**Figure 3D**). Simulations qualitatively established the effect of this competition on the recoverer fraction (**Figure S10**). Adding cycloheximide^21^, a specific inhibitor of cytosolic ribosome elongation (**Figure S11A**), with chloramphenicol blunted the effect of chloramphenicol (**Figure 5C**), consistent with competition between cytosolic and mitochondrial ribosomes controlling metabolic bistability.

**Figure 5.**
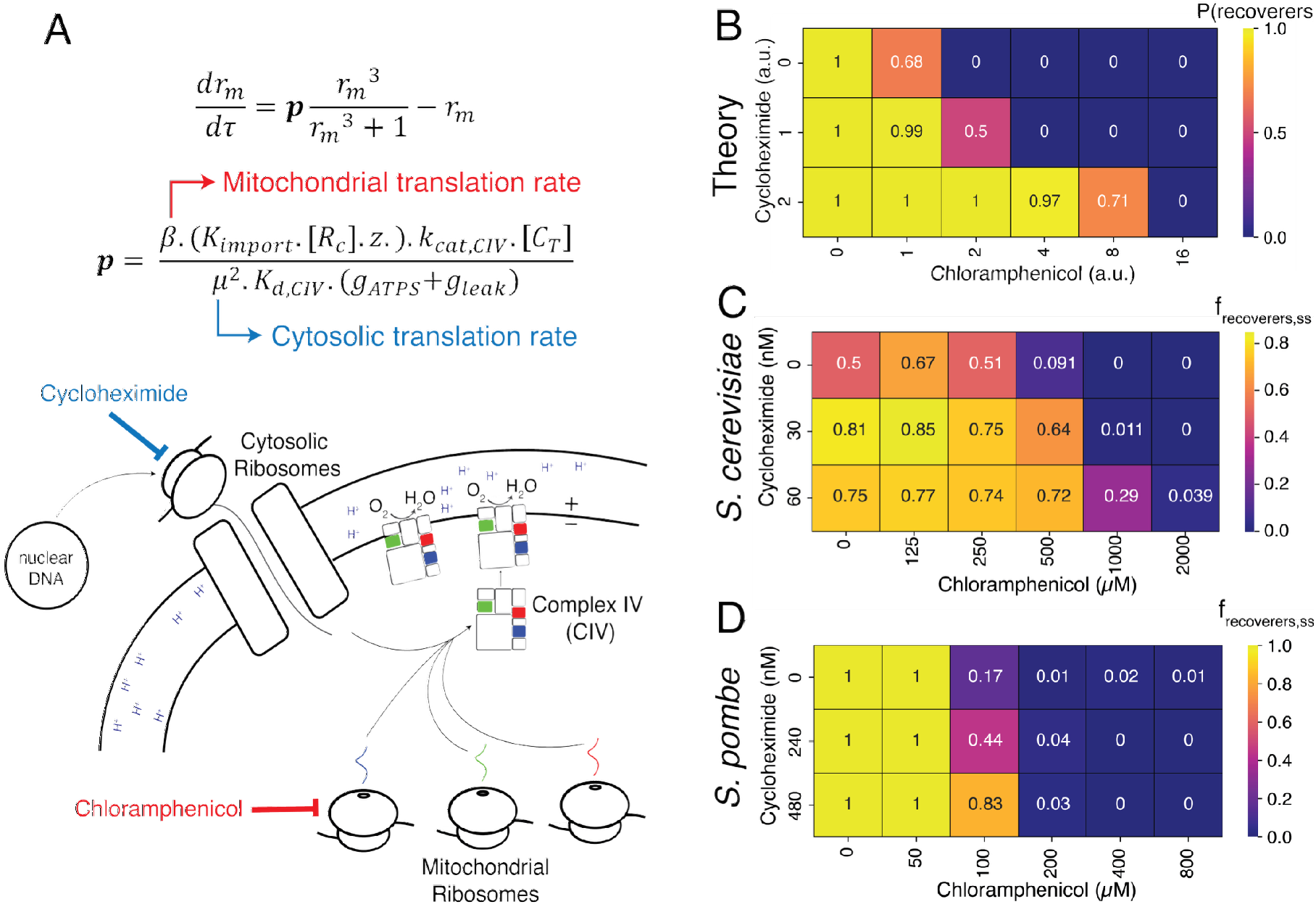
Modulating relative cytosolic and mitochondrial translation rates regulates the balance of arrestors and recoverers (A) A quantitative biophysical model predicts global antagonism between mitochondrial and cytosolic translation. (B) Prediction of changes in the recoverer fraction on combinatorial inhibition of cytosolic translation (cycloheximide; arbitrary units) and mitochondrial translation (chloramphenicol; arbitrary units). The P(recoverer), the probability of being a recoverer, was calculated from the stationary distribution as described in Appendix I. (C) Experimental measurement of steady state recoverer fraction in response to combinatorial titration of chloramphenicol and cycloheximide. (D) Experimental measurement of steady state recoverer fraction in *Schizosaccharomyces pombe* in response to combinatorial titration of chloramphenicol and cycloheximide.

### Tuning the global competition between mitochondrial and cytosolic ribosomes reconstitutes bistability in *Schizosaccharomyces pombe*

The molecular components underlying the bistable circuit are conserved from yeast to mammals, including the three mitochondrially encoded subunits of complex IV—a quantitative feature largely conserved across eukaryotes. In the fission yeast, *Schizosaccharomyces pombe*, which diverged from *S. cerevisiae* roughly 400 million years ago, we couldn’t detect a bimodal response to glucose deprivation, consistent with *S. pombe’s* inability to survive without mitochondrial DNA^*22*^. Our model predicts that reducing mitochondrial translation in *S. pombe* should give rise to bistability, generating arrestor and recoverer-like states separated by an unstable intermediate (**Figure 5B**). We therefore asked if we could reconstitute bistability in *S. pombe* using chloramphenicol; at 100 µΜ chloramphenicol *S. pombe* produced a mixture of arrestors and recoverers on transition from glucose to glycerol and ethanol and at this concentration, adding cycloheximide increases the fraction of recoverers (**Figure 5D**). This experiment demonstrates that an evolutionarily conserved competition between cytosolic and mitochondrial ribosomes can produce metabolic bistability.

## Discussion

We have shown that a conserved bistable switch in mitochondrial metabolism locks individual cells in fermenting or respiring states. The fermenting, arrestor, state grows faster on high glucose while the respiring, recoverer, state grows faster on lower glucose concentrations and non-fermentable carbon sources. By visualizing the steps in this circuit, we could predict how individual cells would respond to a worse carbon source based on their metabolic state on glucose. The co-operative assembly of cytochrome oxidase, driven by mitochondrial protein synthesis, provides the non-linearity required for a bistable switch, allowing the two cell states to be produced and maintained at constant external glucose concentrations. The opposing influence of mitochondrial ribosomes and cytosolic ribosomes on the concentration of the respiratory apparatus sets the fraction of arrestors and recoverers.

Stable glucose levels are an exception rather than a rule in environments that range from heterogenous tissues to industrial bioreactors. Unpredictable variations in glucose levels pose a fundamental threat to cellular physiology because of its dependence on ATP production. Fermentation allows faster ATP production^1^ on high glucose supporting fast growth whereas respiration allows more efficient ATP production on low glucose and is required to produce ATP from less favorable carbon sources. Switching from fermentation to respiration requires increased synthesis of respiratory proteins^23^–a slow process. The fast turnover of the cellular ATP pool (seconds) and unpredictable environmental glucose fluctuations (seconds to days) versus the slow retooling of metabolic capacity make bet hedging, the production of alternative physiologies within a genetically identical population, such as arrestors and recoverers, an attractive solution. A bistable switch enables a population of cells to stochastically switch between respiration and fermentation, maintaining a metabolic portfolio that guards against unpredictable environments. Because the steady-state fraction of the two states is sensitive to the glucose levels, a population can re-adjust its portfolio based on the current glucose levels.

The Warburg effect, fermentation in the presence of oxygen, is a hallmark of cancer cells^24^ and has been proposed to allow faster cell growth and proliferation. Because cancer cells possess all the components of the bistable circuit in yeast, we speculate that their metabolic rewiring could be explained by the competition between cytosolic and mitochondrial protein synthesis we report in this study. Fast growth driven by oncogenic transformation forces mitochondria to also divide rapidly^25^. Rapidly dividing embryonic stem cells also display phenotypes that range from a mixture of fermentation and respiration to pure fermentation^26^. In both cases, we argue fast mitochondrial growth, driven by cytoplasmic protein synthesis, dilutes the respiratory machinery and reduces respiratory ATP production providing a mechanistic basis for the Warburg effect.

## Acknowledgements

The authors thank Jane Kondev, Timothy Mitchison, Allon Klein and Matthew Vander Heiden for feedback on the results. The authors also thank Michael T. Laub, Pankaj Mehta and Michael M. Desai for critically reading the manuscript and providing feedback. We want to thank members of the Bauer flow cytometry core and the mass spectrometry core at Harvard University. We also thank Danesh Moazed for providing the efflux pump deficient *S. pombe* strains. Figure S1 was partly created with BioRender.com. Claude (Anthropic) and Perplexity were used to generate code from data analysis and figure generation. The code was verified through manual inspection and control tests.

## Author Contributions

P.N. and A.W.M. conceptualized the study. P.N. performed experiments and analyses. P.N. and A.W.M. wrote the manuscript.

## Data, code, and materials availability

Scripts and processed data required for reproducing all the figures are available at: https://github.com/piyushnanda-sysbio/mitochondrial_bistability. Scripts along with the raw data are available at Zenodo: https://doi.org/10.5281/zenodo.22019404. Request for strains or plasmids created in the study should be addressed to the corresponding authors.

## Supplementary Materials

Materials and Methods

Appendix I

Figures S1 to S11

Tables S1 and S2

Movies S1-S3

**Main Figures**

