## Supplementary Materials for "Translational antagonism enables bet-hedging in yeast via mitochondrial bistability"

### Materials and Methods

#### Cell culture and molecular biology

*S. cerevisiae* W303 *(MAT***a***)* was grown in synthetic media containing complete supplemented media (0.7g/L), yeast nitrogen base with ammonium sulphate (6.7 g/L) and the specified carbon source at 30˚C. Cultures were maintained at early exponential phase (<3e7 cells per ml), by constant dilution, for all experiments, to keep glucose levels constant in the media and grown in liquid culture for at least 24 hours from a single colony. The concentration of ethanol and galactose were set at 1% (v/v) and 2% (w/v) in the post-shift media. The concentration of ethanol was chosen such that its toxic effects, which occur beyond 2% (data not shown), are minimized.

*S. pombe h+* was grown in YES media (0.5% (w/v) Yeast Extract supplemented with 50 mg/L of each of alanine, lysine, histidine, leucine and uracil). Glucose was used as a carbon source at 3% (w/v). For post-shift media, 1% (v/v) glycerol with 2% (v/v) ethanol was used as a mixed carbon source.

For treatment with inhibitors, cells were pre-grown in the appropriate pre-shift media condition for 12-16 hours before being shifted to various concentrations of the inhibitors. The cells were then grown for at least 16 hours in the presence of inhibitors for the recoverer fraction to reach steady state. No further change in recoverer fraction with time confirmed a dynamic equilibrium for all the concentrations. For washing out inhibitors or for up/down shifts, the cells were spun in a 96-well hydrophilic filter plate (AcroPrep plates with 0.45 μm Supor membrane) at 1800 RPM for 10 s. The cells were immediately suspended in the post-shift media (with a new concentration of the inhibitor) and washed twice in the new media to remove any residual drugs. Media with drugs were prepared freshly before experiments. For chloramphenicol titration experiments in *S. cerevisiae* the gene encoding Pdr5p (a multidrug efflux pump) was deleted to increase drug sensitivity. For *S. pombe,* a strain harboring deletion of the genes encoding the two major drug efflux pumps, Bfr1p and Pmd1p, was used.

Cloning was performed either by Gibson assembly or Golden Gate assembly. Constructs were integrated into the genome either using integrative vectors (for large constructs), or using CRISPR Cas9, with standard LiAc/PEG transformation. Donor and constructs were codon optimized for *S. cerevisiae.* Integration or editing was confirmed by colony PCR and Sanger sequencing.

To measure the assembly of cytochrome oxidase using Split-FAST, Cox4 was tagged at the C-terminal with NFAST and Cox5a was tagged at the N-terminal with CFAST10 with an intervening GGGGGSGGGGGS linker. A single CRISPR Cas9 plasmid based on Multiplex Yeast Toolkit (MYT) was used to simultaneously target both the loci and donors with ~200 bp of sequence homology on either side of the integration site were used.

#### Quantitative Fluorescence Microscopy

Fluorescence intensity was measured using a Spinning Disk (Yokogawa) microscope and appropriate laser lines. A 60X, 1.4 NA Objective (Nikon) was used for all experiments. Z-Stacks with 0.4 μm between successive planes were collected and were set to include entire volume of yeast cells. Laser power (minimized for long-term timelapse imaging), gain and exposure time were kept fixed for all comparative analysis. All images used for comparative analysis were taken in a single round of imaging using automated stage control and multi-dimensional imaging program in Metamorph software (Molecular Devices). Cells were loaded into 384 well plates by spinning down a dilute culture of cells at 300 g for 1 min and immediately imaged. Fluorescence intensity was normalized to a constitutively expressed marker in the specific cellular location to account for focal drift or non-uniformity in field of view.

**High-throughput estimation of recoverer fraction and single-cell lag phase (Figure S1)**

After steady state growth in appropriate media, cells were shifted to either 1% ethanol or 2% galactose. Briefly, cells were spun down at 1800 RPM for 10 s through a 96 well hydrophilic 0.45 µm Supor membrane (Acroprep) attached to a 96 well deep-well plate to receive the flow-through. The cells were transferred to ethanol or galactose by repeating the steps above with fresh media thrice. The cells were diluted to appropriate cell density to reduce crowding in the field of view. For long-term tracking of cells, 384 well glass bottom plates were coated in 2 mg/ml Concanavalin A (with 5 mM manganese chloride and 5 mM calcium chloride) for 5 mins at room temperature, followed by washing out any unbound Concanavalin A with sterile MilliQ water. About 40 µL of cell culture at appropriate cell density was spun down to the bottom of the plate at 300 g for 1 min. The cells were incubated at 30 deg C during imaging.

Cells were imaged every 30 mins for 36 hours to estimate the fraction of recoverers and their single cell lag times. At least three fields of view were imaged per replicate per condition. The cell density was adjusted to make sure the cells are well separated by keeping the optical density (OD600) below 0.01 or 3E5 cells per ml.

**CellASIC microfluidic growth**

CellASIC Y04C chips were used with the programmable perfusion set-up for tracking single cells during the metabolic shifts. Briefly, all the channels (1-6) were activated for 5 mins at 2 psi to make sure the connecting channels between the reservoir well and the imaging chamber did not have residual storage buffer. This is important for fast starvation experiments where the cells would otherwise encounter storage media prior to their exposure to the post-shift media, triggering a premature starvation response. The imaging chamber was coated with 2 mg/ml Concanavalin A (with 5 mM Manganese chloride and 5 mM Calcium chloride) for 5 mins at 2 psi, followed by 1X PBS for 5 mins at 2 psi to washout unbound Concanavalin A. The pre-shift media was perfused through the chamber for 5 mins at 2 psi before loading the cells. The cells were loaded into the CellASIC chamber at 4 psi-8 psi for <20 s.

Cells were grown in pre-shift media for 1-4 hours by perfusing the synthetic media at 2 psi. Cells were shifted to the post-shift media by perfusing the post-shift media at 4 psi for 5 mins to rapidly washout any glucose from pre-shift media. The perfusing rate of the post-shift media was set back to 2 psi for 8-12 hours. Dextran Texas-Red (40,000 Da) was used to calibrate the residence time of the media and for calibrating the exchange time (time taken for shifting from one media to the another) with the CellASIC ONIX software media switch indicators.

**Quantification of ATP:ADP with PercevalHR**

We used a yeast codon optimized PercevalHR under the strong *TDH3* promoter that was integrated in the yeast genome. The PercevalHR signal is affected by the pH making it necessary to simultaneously measure the cytosolic pH, which also changes rapidly like the ATP:ADP ratio. A second strain expressing SEP (super ecliptic pHluorin, non-ratiometric) fused to mRuby2 was used to simultaneously track pH in the same chamber. The mRuby2 was used both for ratiometric pH imaging and for differentiating it from the strain expressing PercevalHR. The SEP and the PercevalHR strains were grown separately for 24 hours in CSM+2% Glucose, mixed in 1:1 ratio and loaded into the CellASIC chamber as described above.

The mock shift was performed by shifting the media from CSM+2% Glucose to CSM+2% Glucose using the fluidics parameters stated above. Cells were imaged at various points during the shift by manually adjusting the imaging frequency in the multi-dimensional acquisition feature of the MetaMorph software (Molecular Devices). The exact same fluidics and imaging parameters were used as for the CSM+2% Glucose to CSM+2% Galactose shift.

At the end of the experiment, the pH influence on PercevalHR was measured by the pre-pulse method as described earlier, in both glucose replete and glucose depleted conditions. Briefly, cells were exposed to 1 mM ammonium chloride in CSM+2 % Glucose or CSM + 2% Galactose for 5 mins at 2 psi. The pH of the cytosol has a non-monotonic response due to consecutive alkalinization, and acidification of the cytosol induced by external ammonium chloride. The signal from the PercevalHR and pH sensor was recorded. The PercevalHR signal was regressed against the pH signal. The slope was used to correct for the effect of pH on PercevalHR in arrestors and recoverers. The cells were assigned as arrestors and recoverers based on their growth recovery on CSM + 2% Galactose.

**Biochemical measurement of ATP:ADP**

To measure the absolute ATP:ADP ratio before and after glucose deprivation, cells growing at steady state in 2% glucose and after 2 hours following abrupt shift to 2% galactose, were sprayed directly into pure methanol maintained at -40 deg C using ethylene glycol and ethanol dry-ice bath. The pellet was collected by spinning down the mixture at 3000 RPM for 2 mins in a precooled (4 deg C) centrifuge. The pellet was resuspended in hot ethanol buffered with 10 mM HEPES at pH 7.2 and the mixture was shaken at 80 deg C for 3 mins. The supernatant was collected by spinning down the mixture at 20,000 g for 2 mins and evaporated under vacuum (SpeedVac) at 30 deg C for 1.5 hours. The residual material was resuspended in HPLC grade water. The ATP:ADP ratio was estimated using ATP:ADP ratio assay kit from Sigma Aldrich following manufacturer's instructions.

**Measurement of mitochondrial membrane potential**

100 nM TMRM was directly added to the growth media from a 100 µM TMRM stock in DMSO. Cells were grown for 30 mins in 100 nM TMRM before being loaded into a CellASIC chip. TMRM was also added to the media in the CellASIC chip at 100 nM concentration for live imaging experiments. Measurements were performed only when TMRM reached a steady-state concentration in the chip, which in the case of high glucose took >2 hours.

**FACS sorting of FBP reporter strains**

A BD FACS Aria I was used for sorting cells based on the signal from the FBP reporter. Briefly, a culture of exponential grown cells in CSM+0.05% glucose was loaded into the input port maintained at 20 deg C and spinning at maximum speed. The top 15% and bottom 15% fractions were directly sorted into collection tubes with CSM+0.05% glucose maintained at 20 deg C. A 70 µm nozzle was used for sorting the yeast cells and 'purity' mode was used. Roughly 400,000 cells were sorted for each of the fractions. The cells were plated to confirm the number of viable cells (colony forming units, CFUs).

For determining the optimal gates, the signal from the fluorescence channel (FITC) was plotted against the cell size (FSC-A). Two quadrilateral gates (Top and bottom 15%) were chosen such that the cell size distributions overlapped, and the FBP reporter signal distributions had limited overlap. This allowed for sorting cells based on their FBP signal rather than their cell size or morphology. The efficiency of the sorting was estimated by measuring the fraction of recoverers in the two fractions by shifting them to ethanol and galactose as described previously. A strain lacking the CggR R250A TF was used as a negative control.

The sorted cells were immediately used for ethanol production rate and the oxygen consumption rate assays. For the assays, the cells were concentrated by using a hydrophilic filtration unit to reach the cell density necessary for sensitive measurements in the assays.

**Ethanol production rate measurement**

An Amplite™ Ethanol Quantification Kit was used for measuring ethanol concentration in the media. Briefly, supernatant from the culture was collected by centrifuging at maximum speed. Samples were collected at t=0 and t=30 mins after sorting and the ethanol concentration in the media was measured following the manufacturer's instructions. The number of cells per ml of the media, estimated by counting CFUs, was used to calculate ethanol production rate for the two fractions.

**Oxygen uptake rate measurement**

A modified protocol using OxoPlates (PreSens Precision Sensing GmbH) was used to measure oxygen uptake rate in the sorted fractions. The sorted cultures were concentrated and transferred to an OxoPlate. The plate was immediately sealed using a gas impermeable film and the instantaneous rate of oxygen depletion was measured using the time-resolved fluorescence mode. The plates were calibrated using air saturated water or 1% sodium sulfite solution for maximum oxygen tension and zero oxygen respectively. The measurements were performed within 1 hr to minimize interference due to cell growth in the chamber. Absolute oxygen uptake rate was calculated using the cell density and the rate of oxygen depletion.

**Quantification and data analysis**

**Cell segmentation and quantification**

Masks of high focus brightfield images of the cells were obtained using Cellpose cyto3 model with custom scripts written in Python. Fluorescence quantification of single-plane images was performed by subtracting the background fluorescence from all the cells and averaging over the region within the mask periphery. For fluorescence quantification of multi-plane images (mitochondria), the mitochondrial region was segmented using an independent fluorescent marker. The sum intensity z-projection of the 3D stack was used to calculate the mean intensity within the mitochondria periphery.

The masks generated from Cellpose were used to manually annotate the cells as arrestors or recoverers using Napari based on growth recovery following shift to galactose. Fluorescence quantification was performed only for the last frame before carbon source switch.

**Estimating of recoverer fraction from brightfield timelapse movies**

A custom pipeline was developed to estimate recoverer fraction and their single cell lag times from brightfield low magnification timelapse movies. The data was obtained as described above by imaging

Concanavalin A immobilized cells in a 384 well plate for 36 hours. Images were obtained every 30 mins. The fraction of recoverers was defined as the fraction of cells that underwent at least one complete cell division within 36 hours. The brightfield images were segmented using Cellpose cyto3 model. A modified Hungarian algorithm was used to perform cell-tracking with cell division. Briefly, the distance matrix between consecutive frames was obtained from the centroid of the masks. The entries of the distance matrix were duplicated to allow mapping between (i) the current and next locations of the mother cell (ii) the mother and the daughter cell formed in the next frame. Because the cells are sparsely located in the field of view, the distance could be used to assign mother-daughter links. For speeding up the tracking algorithm, the image was sliced into overlapping frames to constrain the assignment problem. The penalties of the assignment problem were calibrated to reproduce manually counted recoverer fractions. For each cell, a lineage tree was obtained that connects the masks at time t to its future state. The number of new branches, representing formation of daughter cells, was used to declare a lineage that showed growth recovery and cell division. The single cell lag time was calculated by the time taken for a new branch to arise. The recoverer fraction was calculated by the fraction of lineages that produced at least one branch in 36 hours.

### 1 Appendix I: Ordinary Differential Equation and Probabilistic Models of the Bistable Switch

#### 1.1 Assembly of complex IV driven by mitochondrial translation

We consider three mtDNA encoded subunits Cox1, Cox2 and Cox3 as $C_1$, $C_2$, and $C_3$ that assemble into a functional complex IV, denoted $C_{\mathrm{IV}}$, along with the nuclear encoded subunits.

Their production dynamics are

$$\frac{dC_i}{dt} = \beta R_m - \mu C_i, \quad i = 1, 2, 3. \tag{1}$$

At steady state $\left(\frac{dC_i}{dt} = 0\right)$,

$$C_i = \frac{\beta R_m}{\mu}, \quad i = 1, 2, 3. \tag{2}$$

Let $C_T$ be the average concentration of the nuclear-encoded subunits that form the complex, and let $x = [C_{\mathrm{IV}}]_{\mathrm{eq}}$ be the equilibrium concentration of assembled complex IV. The equilibrium binding relation is

$$K_d^3 = \frac{(C_1 - x)(C_2 - x)(C_3 - x)(C_T - x)}{x}. \tag{3}$$

Assuming $C_i \gg x$, so that $C_i - x \approx C_i$ for $i = 1, 2, 3$, and substituting Eq. (2) into Eq. (3), we obtain

$$K_d^3 = \frac{C_1 C_2 C_3 (C_T - x)}{x} = \left(\frac{\beta R_m}{\mu}\right)^3 \frac{C_T - x}{x}. \tag{4}$$

Rearranging Eq. (4) for $x$ gives

$$x = [C_{\mathrm{IV}}]_{\mathrm{eq}} = C_T \frac{\left(\frac{\beta R_m}{\mu}\right)^3}{\left(\frac{\beta R_m}{\mu}\right)^3 + K_d^3}. \tag{5}$$

#### 1.2 Mitochondrial membrane potential as a function of ETC activity

We assume that ATP synthase and the proton leak behave as ohmic conductors. The mitochondrial membrane potential $\Delta\psi$ is given by:

$$\Delta\psi = \frac{I_{\mathrm{proton}}}{g_{\mathrm{ATPS}} + g_{\mathrm{leak}}}, \tag{6}$$

where $g_{\mathrm{ATPS}}$ and $g_{\mathrm{leak}}$ are the effective conductances of ATP synthase and leak, respectively.

We assume the proton current is proportional to the amount of complex IV:

$$I_{\mathrm{proton}} = k_{\mathrm{cat}} [C_{\mathrm{IV}}]_{\mathrm{eq}}, \tag{7}$$

with turnover rate $k_{\mathrm{cat}}$.

Substituting Eq. (7) into Eq. (6), we obtain

$$\Delta\psi = \frac{k_{\text{cat}}[C_{\text{IV}}]_{\text{eq}}}{g_{\text{ATPS}} + g_{\text{leak}}}. \tag{8}$$

Using Eq. (5) for $[C_{\text{IV}}]_{\text{eq}}$, we get

$$\Delta\psi = \frac{k_{\text{cat}} C_T}{g_{\text{ATPS}} + g_{\text{leak}}} \cdot \frac{\left(\frac{\beta R_m}{\mu}\right)^3}{\left(\frac{\beta R_m}{\mu}\right)^3 + K_d^3}. \tag{9}$$

##### 1.3 Mitoribosome flux into mitochondria as a function of $\Delta\psi$

Mitochondrial ribosome dynamics are modeled as

$$\frac{dR_m}{dt} = K_{\text{import}}\, z\, \Delta\psi\, [R_c] - \mu R_m, \tag{10}$$

where $K_{\text{import}}$ is the import rate constant, $z$ is an effective charge factor, $\Delta\psi$ is the membrane potential, and $[R_c]$ is the cytosolic ribosome pool.

Substituting Eq. (9) into Eq. (10) yields

$$\frac{dR_m}{dt} = K_{\text{import}}[R_c]\, z\, \frac{k_{\text{cat}} C_T}{g_{\text{ATPS}} + g_{\text{leak}}} \cdot \frac{\left(\frac{\beta R_m}{\mu}\right)^3}{\left(\frac{\beta R_m}{\mu}\right)^3 + K_d^3} - \mu R_m. \tag{11}$$

This equation closes the feedback loop between mitoribosome abundance, complex IV, and $\Delta\psi$.

##### 1.4 Non-dimensional form of the equation

We nondimensionalize the system to highlight the switch-like response. Define the dimensionless mitoribosome concentration:

$$r_m = \frac{\beta R_m}{\mu K_d}, \tag{12}$$

such that the nonlinear term takes the form $r_m^3/(r_m^3 + 1)$.

Define dimensionless time rescaled to cell growth timescales, $\tau = t\mu$.

Collecting constants into a single parameter $p$, Eq. (11) reduces to

$$\frac{dr_m}{d\tau} = p\, \frac{r_m^3}{r_m^3 + 1} - r_m. \tag{13}$$

The effective feedback strength parameter $p$ is

$$p = \frac{K_{\text{import}}[R_c]\, z\, \beta\, k_{\text{cat}} C_T}{\mu^2 K_d\, (g_{\text{ATPS}} + g_{\text{leak}})}. \tag{14}$$

A constant basal production term $p_0$, with the same units as p, can be added to (13) that accounts for mitoribosomal precursor entry that is independent of the membrane potential,

$$\frac{dr_m}{d\tau} = p_0 + p\,\frac{r_m^3}{r_m^3+1} - r_m. \tag{15}$$

Equation (15) captures the core switch-like behavior: for appropriate $p$, the cubic nonlinearity in $r_m^3/(r_m^3+1)$ can generate sharp transitions between low and high $r_m$ steady states, corresponding to low and high mitochondrial translation regimes.

##### 1.5 Simulating the stochastic behaviour of the bistable switch

From (15), we extract the production term and the dilution term to model a 1D birth-death process,

Let $n$ be the number of mitoribosomes and $\Omega$ be the system size such that $\mathrm{r}_m = \frac{n}{\Omega}$.

We define the production term as,

$$g(r_m) = g(\frac{n}{\Omega}) = p_0 + p\,\frac{r_m^3}{r_m^3+1} \tag{16}$$

The dilution term is,

$$d(\frac{n}{\Omega}) = r_m \tag{17}$$

With $P(n)$ as the probability of having $n$ mito-ribosomes, the system will satisfy the detailed balance $P(n)g(\frac{n}{\Omega}) = P(n+1)d(\frac{n+1}{\Omega})$.

Recursively solving, we get:

$$P(n) = P(0)\prod_{k=1}^{n}\frac{\Omega \cdot g(\frac{k-1}{\Omega})}{\Omega \cdot d(\frac{k}{\Omega})} \tag{18}$$

Because $d(\frac{n}{\Omega})\ = r_m$, the denominator is n $(= \Omega \cdot r_m)$, we get:

$$P(n) = P(0)\prod_{k=1}^{n}\frac{\Omega \cdot g(\frac{k-1}{\Omega})}{k} \tag{19}$$

Taking the logarithm on both sides to simplify, we get:

$$ln(P(n)) = ln(P(0)) + \sum_{k=1}^{n}[ln(\Omega \cdot g(\frac{k-1}{\Omega})) - ln(k)] \tag{20}$$

We can recursively solve this equation to obtain a series of ln(P) values corresponding to $n$, ranging from 0 to $n_{max}$. Because $\frac{r_m^3}{r_m^3+1} < 1$, the upper stable fixed point satisfies $r_m < p_0 + p$. Therefore, we set $n_{max} = (p_0 + p + 5) \cdot \Omega$ to sample the tail of the distribution beyond the upper stable fixed point.

We recover the probability, $P(n)$, from the $ln(P(n))$ by exponentiating it: $P(n) = e^{ln(P(n))}$

We normalize $P(n)$:

$$P_{norm}(n) = \frac{P(n)}{\sum_n P(n)} \tag{21}$$

This gives a distribution of $P_{norm}(n)$ corresponding to the probability of finding $n$ mitoribosomes.

From the differential equation, we calculate the unstable fixed point, $r_{m,unstable}$ and find $n_{unstable} = \Omega \cdot r_{m,unstable}$.

$n_{unstable}$ corresponds to the number of mito-ribosomes at the unstable fixed point.

Using $n_{unstable}$, we calculate the probability of recoverers, $P(recoverers)$ as:

$$P(recoverers) = \sum_{n>n_{unstable}} P_{norm}(n) \tag{22}$$

#### Supplementary Figures

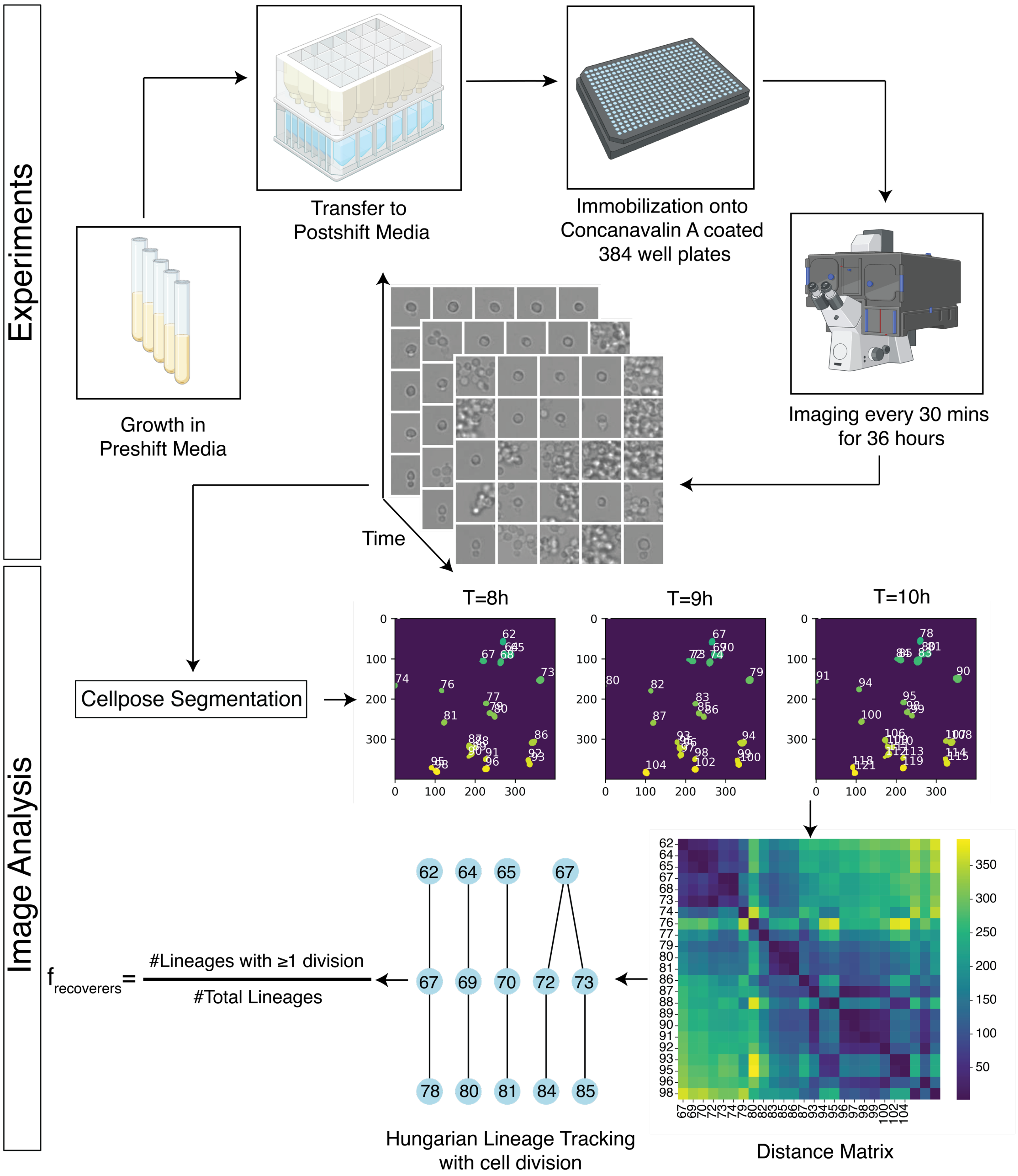


**Figure S1** Experimental workflow for measurement of recoverer fraction by timelapse microscopy and automated image analysis pipeline for estimation of recoverer fraction. Further details in experimental procedures.

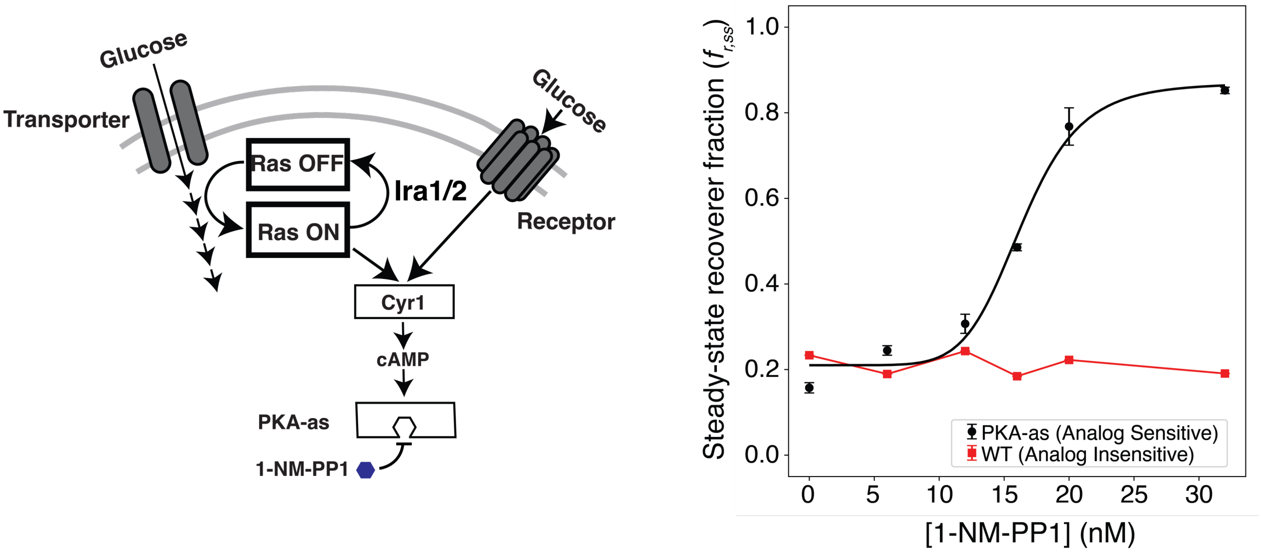


**Figure S2** Monotonic increase in the steady state recoverer fraction on reducing the activity of cAMP-dependent protein kinase A. A strain with analog sensitive alleles of the genes for the three paralogous subunits of the catalytic cAMP-dependent protein kinase (*tpk1-as*, *tpk2-as* and *tpk3-as*) was used to titrate the pathway activity using the Shokat inhibitor 1-NM-PP1. The graph shows the steady state recoverer fraction of cultures growing on 2% glucose that were treated with the indicated 1-NM-PP1 concentrations. A wild type strain without the analog sensitive alleles was used as a negative control.

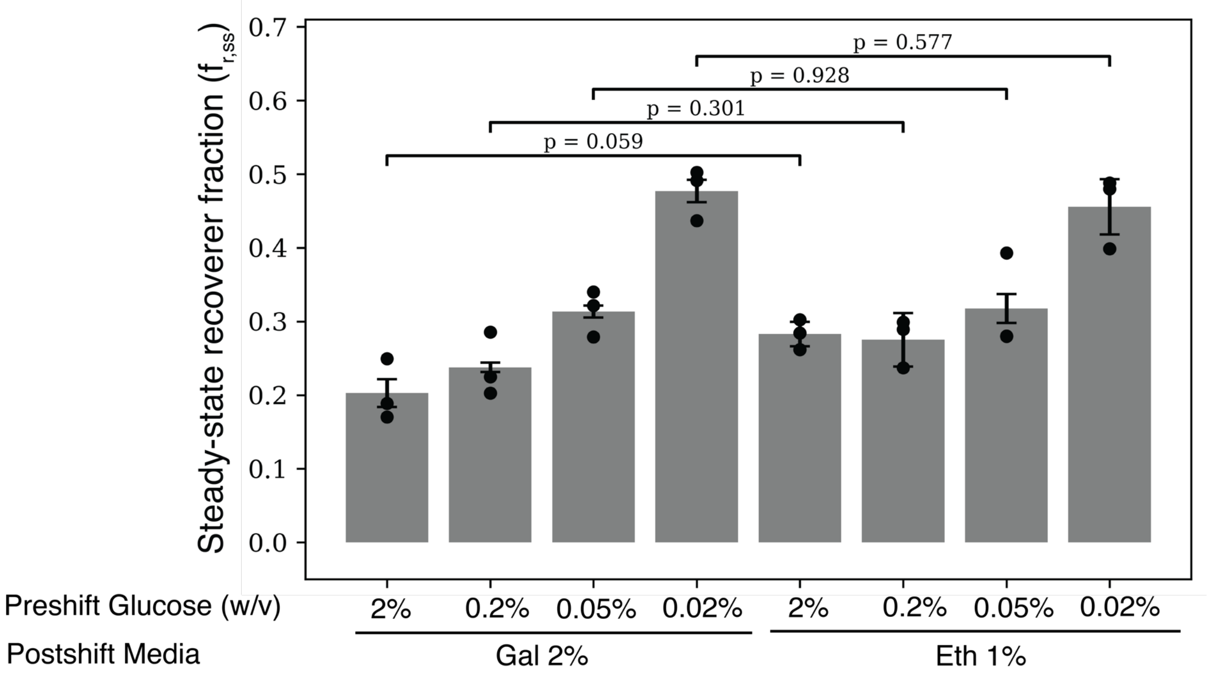


**Figure S3** The steady state recoverer fraction is insensitive to the post-shift carbon source identity and depends on pre-shift glucose concentration. Cells were grown on the indicated concentrations of glucose and then shifted to 2% galactose or 1% ethanol. Welch's t-test was used to compute the statistical significance for the pairwise comparisons shown in the figure.

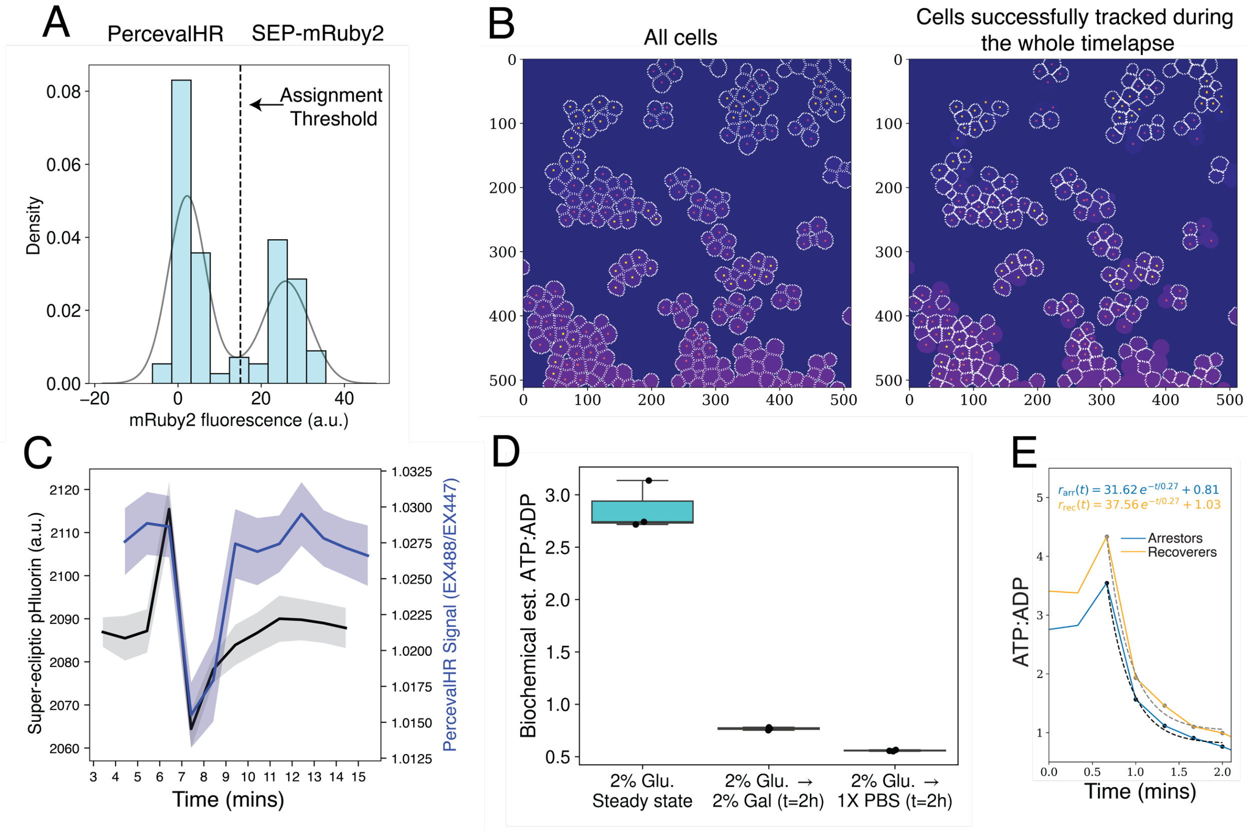


**Figure S4** Image analysis and biochemical calibration of PercevalHR (A) A strain expressing PercevalHR was pooled with a strain expressing the pH sensor, super-ecliptic pHluorin-mRuby2 (SEP-mRuby2) construct. The mRuby2 channel was used to distinguish the two strains based on its higher signal in cells expressing the super-ecliptic pHluorin sensor. (B) A fully automated pipeline was used to track single cells for the full duration of the ATP:ADP and pH measurement experiment. The right image shows the fraction of cells that we successfully tracked for the whole duration by white dotted boundaries. The left image shows all the cells. (C) Pre-pulse method for measuring pH sensitivity of PercevalHR by subjecting the two strains to 1 mM ammonium chloride, which briefly alkalinizes the cytosol followed by acidification. The correlation between the two signals was used to subtract the contribution of pH towards changes in PercevalHR signal. (D) ATP:ADP was biochemically estimated using the Luciferase assay on cells adapted to 2% glucose and 2 hours following transition to 2% galactose. A shift was also performed to 1X PBS (pH 7.2) to show that the assay can detect lower ATP:ADP than measured after 2 h in 2% galactose. (E) The initial dynamics of ATP:ADP in arrestors and recoverers was fit to an exponential decay function to obtain the time constant and the ATP:ADP at the minima.

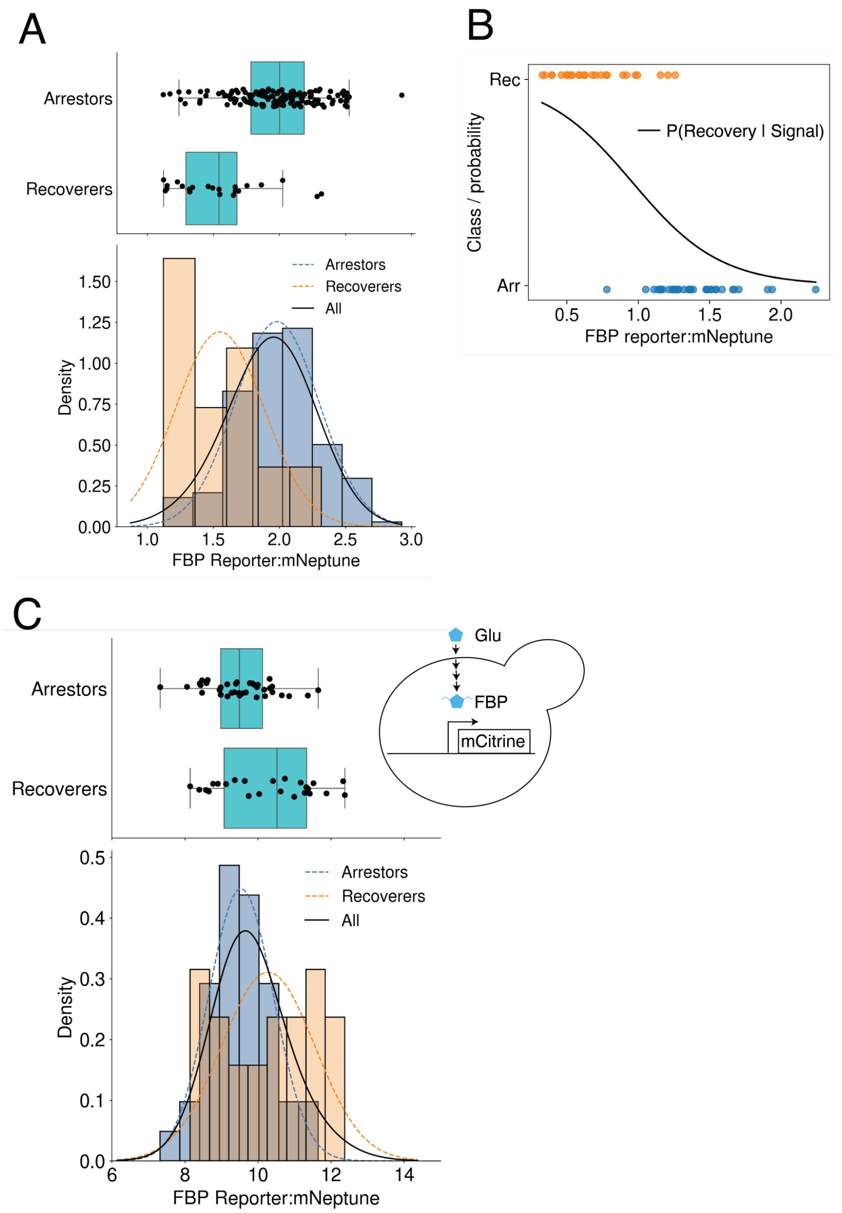


**Figure S5** Measurement of FBP signal in arrestors and recoverers (A) Distribution of FBP signal in arrestors and recoverers, grown on 2% glucose, normalized to mNeptune constitutively expressed from the *ACT1* promoter. (B) Logistic regression of FBP signal from cells growing on 0.05% glucose to calculate a classification metric, the Matthews Correlation Coefficient. (C) Negative control experiment for the FBP sensor in a strain that lacks the CggR transcriptional repressor producing constitutive reporter levels in both arrestors and recoverers. Arrestors and recoverers states were assigned based on their recovery after shift to 2% galactose.

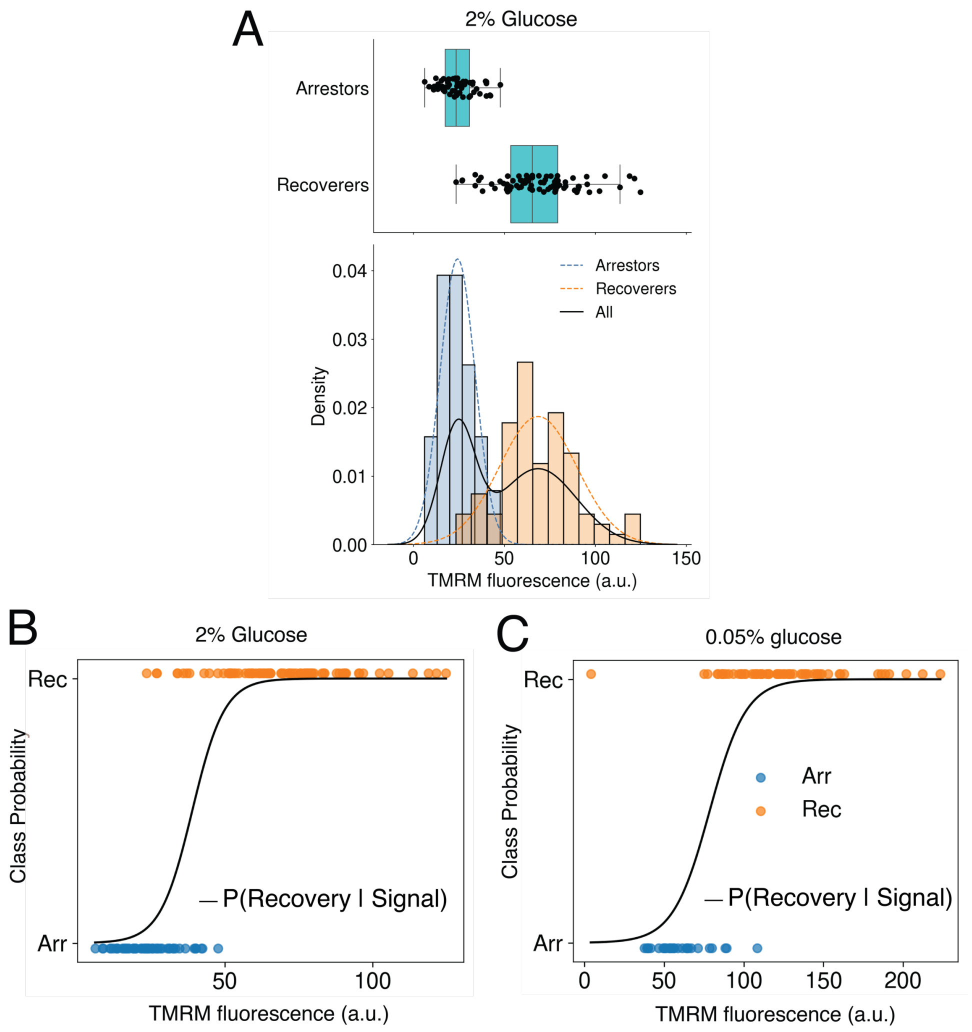


**Figure S6** Logistic regression of TMRM signal during glucose growth to calculate a classification metric, the Matthews Correlation Coefficient (MCC) (A) Distribution of TMRM signal in arrestors and recoverers during growth on 2% glucose. (B) and (C) Logistic regression to calculate classification metrics for growth on 2% glucose and 0.05% glucose respectively.

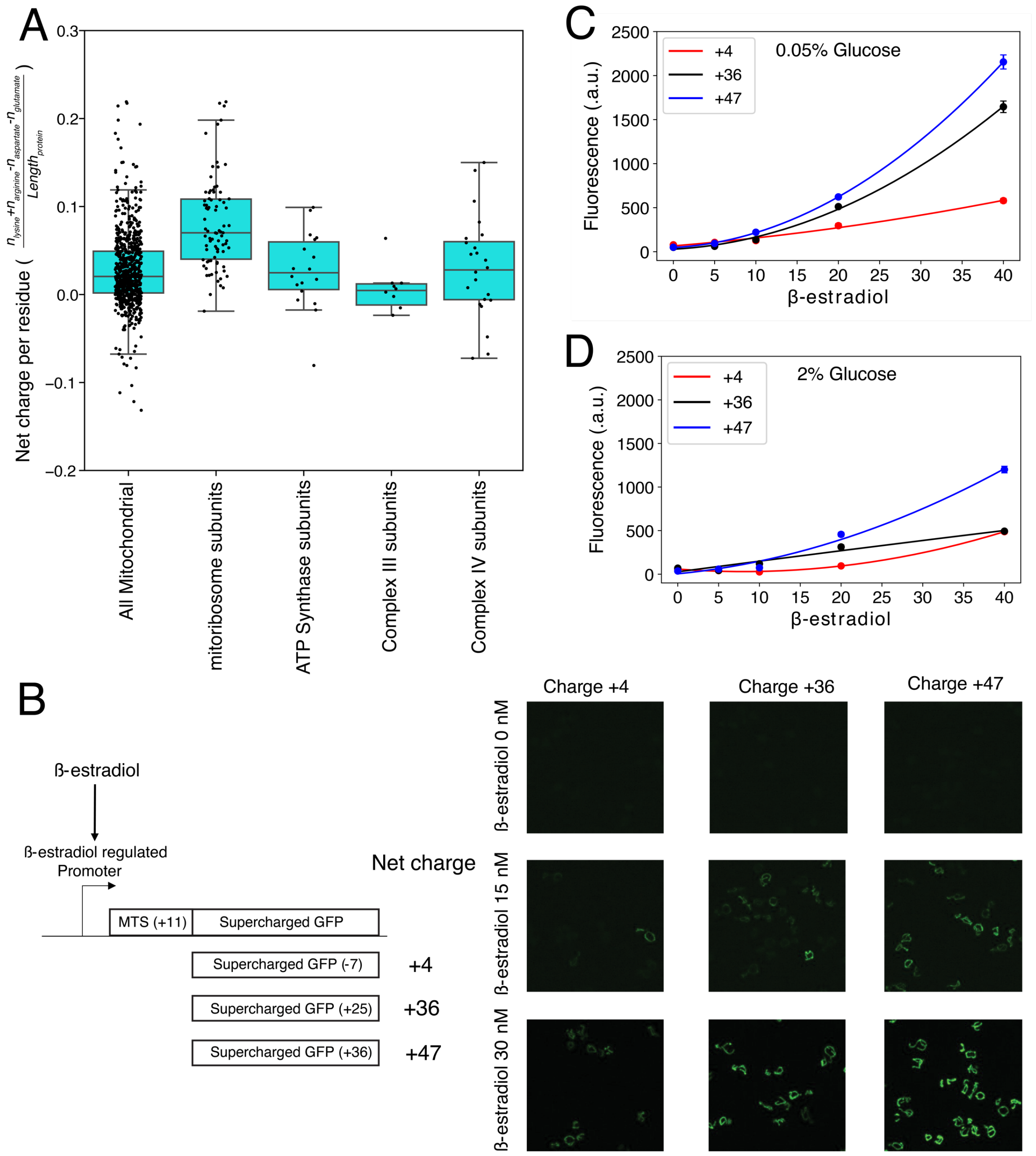


**Figure S7** Differential charge produces membrane-potential dependent import of mitochondrial targeted proteins (A) Polypeptide net charge density for all nuclear-encoded mitochondrially imported polypeptides that participate in the complexes of the electron transport chain, ATP synthase, and mitochondrial ribosomes. (B) An assay based on inducing supercharged GFP production to measure differential import due to differential charge density. MTS is the mitochondrial transit sequence. Quantification of fluorescence signal from mitochondria for various supercharged GFP proteins (charge +4, +36, and +47) expressed at different levels at 0.05% glucose (C) and 2% glucose (D) resulting in higher and lower mitochondrial membrane potential respectively.

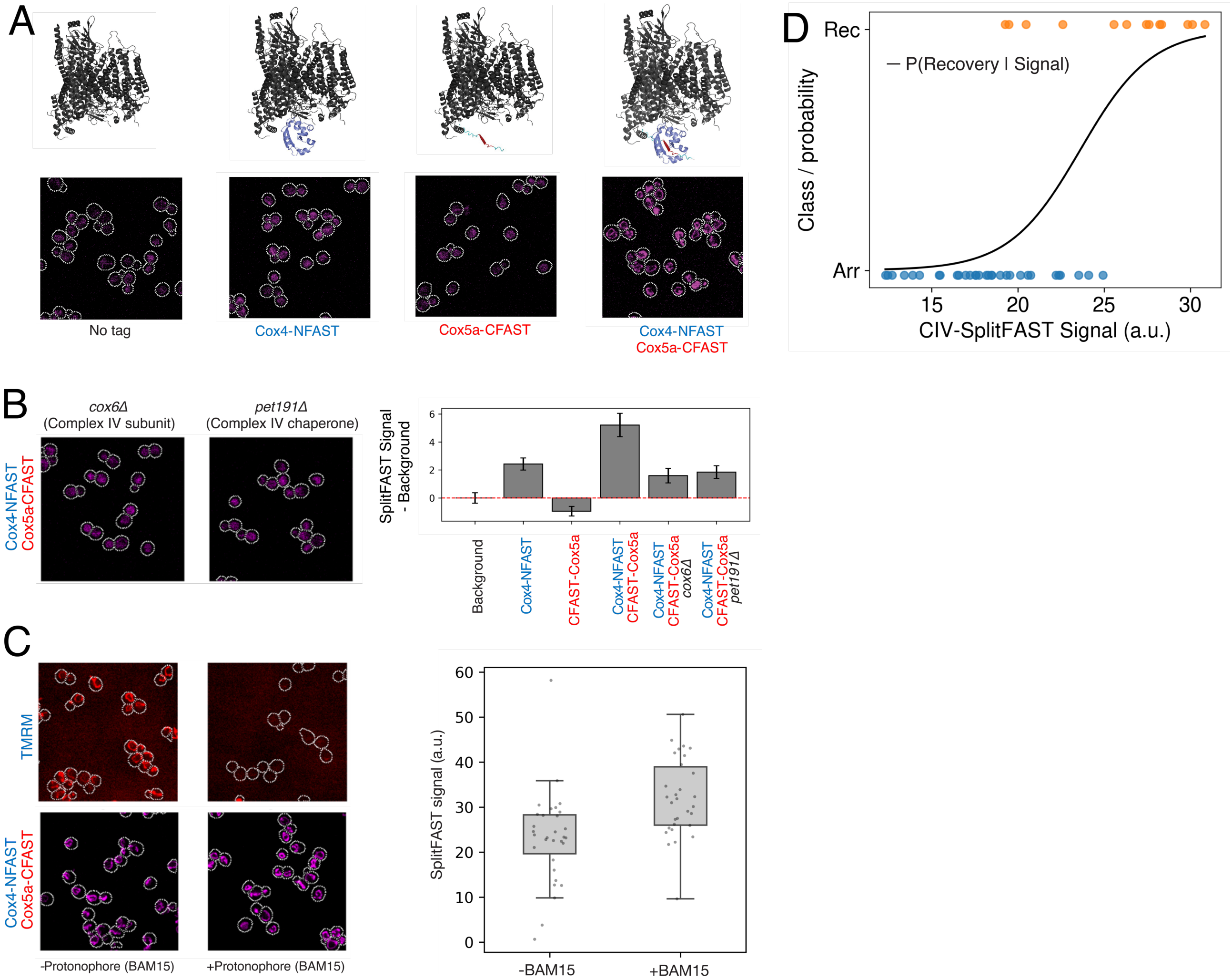


**Figure S8** The complex IV assembly reporter specifically reports the concentration of the assembled complex and is insensitive to membrane potential (A) Signal from HMBR (fluorogen) bound to splitFAST is maximum when both the CFAST (attached N-terminally to Cox5a) and NFAST (attached C-terminally to Cox4) are present. (B) Split-FAST signal on deletion of structural subunit Cox6 or Complex IV specific chaperone Pet191. (C) Split-FAST signal on addition of mitochondria-specific protonophore BAM15 to estimate effect on fluorogen HMBR uptake. The membrane potential was dissipated before HMBR addition. (D) Logistic regression of signal from the complex IV assembly reporter in arrestors and recoverers for calculation of the classification metric, MCC.

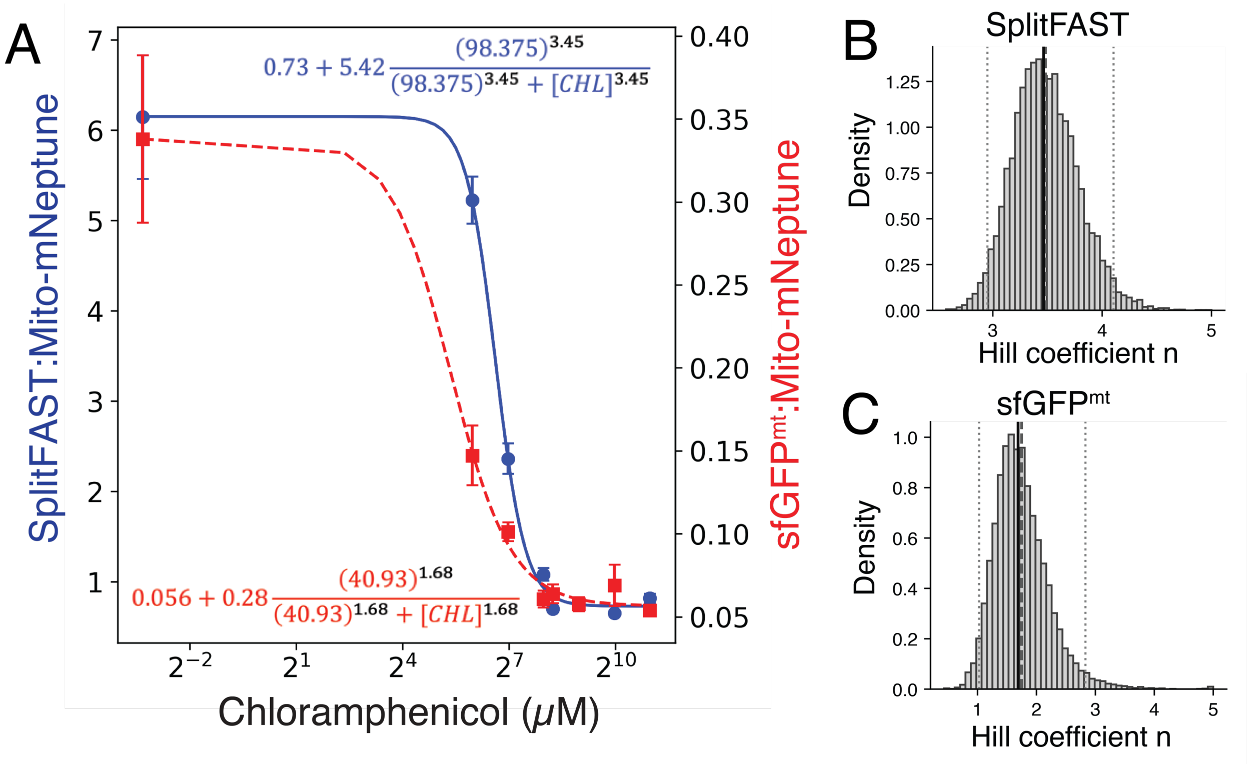


**Figure S9** Measurement of steady-state signal from split-FAST Complex IV reporter and mtDNA encoded sfGFP on chloramphenicol titration in 0.05% (w/v) glucose. (A) A Hill function of the form $y = y0 + y_{max}\frac{k^n}{k^n+[CHL]^n}$ was fit to the data. $n$ is the Hill coefficient and $k$ is the chloramphenicol IC50 in μM. The distribution of Hill coefficients was obtained by sampling the distribution of the residuals between the experimental reporter levels and equation fit for split-FAST signal (B) and the sfGFP$^{mt}$ signal (C).

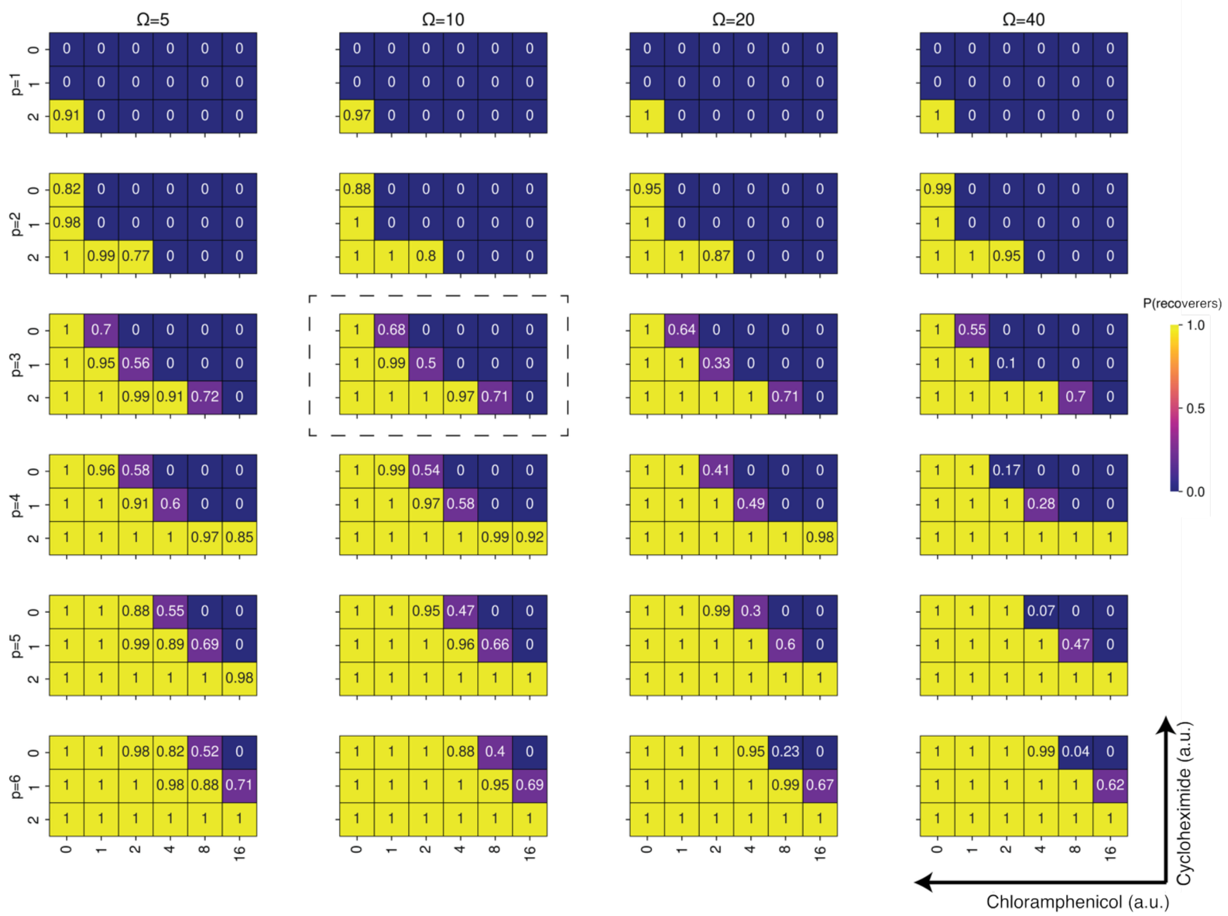


**Figure S10** Stochastic simulation of the bistable switch model Simulated P(recoverers) i.e. Probability of recoverers for various concentrations (arbitrary units) of chloramphenicol (reducing the mitochondrial translation rate parameter) and cycloheximide (reducing the specific growth rate). Briefly, the probabilities were calculated from the stationary distribution of 1D birth-death process that satisfies detailed balance, $P(n+1)d(\frac{n+1}{\Omega}) = P(n)g(\frac{n}{\Omega})$. $d(\frac{n}{\Omega})$ is the dilution rate and $g(\frac{n}{\Omega})$ is the production rate. p is the effective feedback strength parameter (dimensionless) and omega is the system size (volume scale) that converts molecular number to concentrations. The heatmap highlighted by the dashed lines is the one shown in Figure 5B.

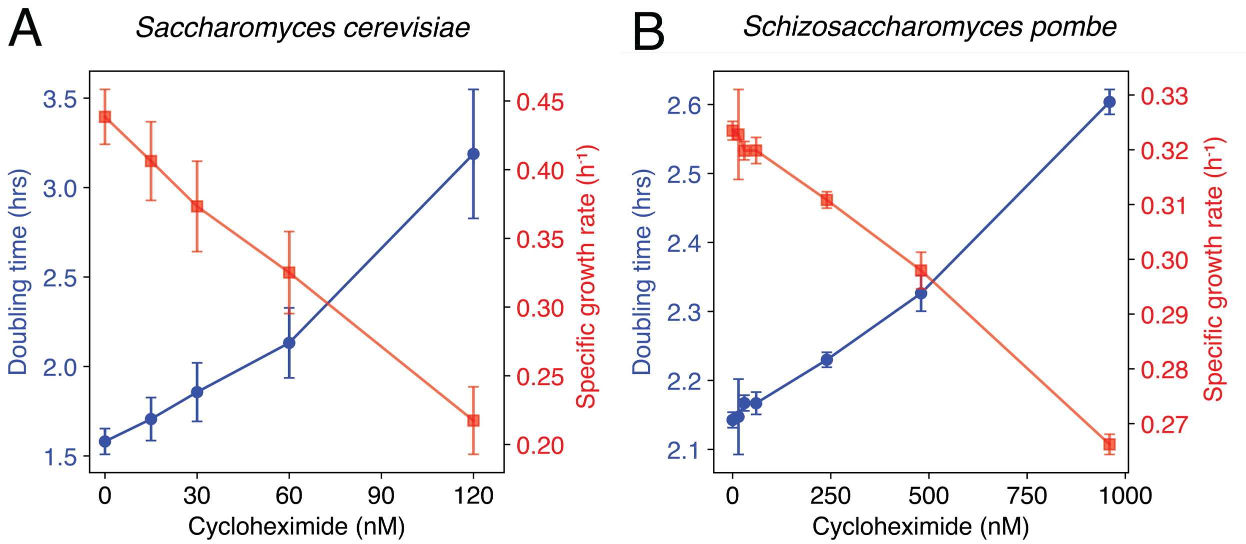


**Figure S11** Measurement of bulk specific growth rate on cycloheximide titration in *S. cerevisiae* and *S. pombe* (A) *S. cerevisiae* cells were grown in 0.05% glucose with cycloheximide for 12 hours and new steady state specific growth rate was measured using a plate reader. (B) *S. pombe* cells were grown in YES + 3% glucose for 12 hours with various concentrations of cycloheximide and their specific growth rate was measured using a plate reader.

**Table S1** *Strains used in the study*

| Strain ID | Strain and genotype | Source |
|---|---|---|
| yPN309 | *S. cerevisiae W303 MATa can1-100 his3^::prACT1-ymCherry-tADH1-His3MX6 BUD4-S288C* | This study |
| yPN315 | *S. cerevisiae W303 MATa can1-100 his3^::prACT1-ymCherry-tADH1-His3MX6 BUD4-S288C ira2^::KanMX4 ira1^::HphMX4* | This study |
| yPN412 | *S. cerevisiae W303 MATa can1-100 his3-11,15 BUD4-S288C his3^::pRS403-pTDH3-yoPercevalHR-tADH1 (pPN143)* | This study |
| yPN698 | *S. cerevisiae W303 MATa can1-100 his3-11,15 BUD4-S288C his3^::pRS403-pTEF1-Super-ecliptic pHluorin-mRuby2-tADH1 (pPN382)* | This study |
| yPN740 | *S. cerevisiae W303 MATa can1-100 ura3-1 trp1-1 tpk1-as tpk2-as tpk3-as pRS403-CGGR-pCGGRO-mCitrine-tADH1 (pPN309) pACT1-mNeptune-ADH1t-LEU2 (pLB118)* | This study |
| yPN742 | *S. cerevisiae W303 MATa can1-100 ura3-1 trp1-1 tpk1-as tpk2-as tpk3-as pRS403-pCGGRO-mCitrine-tADH1 (pPN188) pACT1-mNeptune-ADH1t-LEU2 (pLB118)* | This study |
| yPN585 | *S. cerevisiae W303 MATa can1-100 leu2-3,112 ura3-1 trp1-1 tpk1-as tpk2-as tpk3-as pRS403- pADH1-preSu9-link-mNeonGreen-HIS3* | This study |
| yPN687 | *S. cerevisiae W303 Mat α ade2-1 his3-11,15 trp1-1 leu2-3,112 ura3-1, sfGFPm::cox2, COX2 pADH1-preSu9-link-mNeptune-HIS3 pdr5Δ::KanMX* | Modified *MOY1355* from *Suhm et al. (2018);* |
| yPN735 | *S. cerevisiae W303 can1-100 his3-11,15 ura3^0 BUD4-S288C COX4-2XGGGGGS-NFAST COX5A-MTS-CFAST10-2XGGGGGS-COX5A pACT1-mNeptune-ADH1t-LEU2 pdr5Δ::KanMX* | This study |
| yPN732 | *S. cerevisiae W303 can1-100 his3-11,15 ura3^0 BUD4-S288C COX4-2XGGGGGS-NFAST* | This study |
| yPN733 | *S. cerevisiae W303 can1-100 his3-11,15 ura3^0 BUD4-S288C COX4-2XGGGGGS-NFAST COX5A-MTS-CFAST10-2XGGGGGS-COX5A* | This study |
| yPN734 | *S. cerevisiae W303 can1-100 his3-11,15 ura3^0 BUD4-S288C COX5A-MTS-CFAST10-2XGGGGGS-COX5A* | This study |
| yPN766 | *S. cerevisiae W303 can1-100 his3-11,15 ura3^0 BUD4-S288C COX4-2XGGGGGS-NFAST COX5A-MTS-CFAST10-2XGGGGGS-COX5A cox6Δ::HphMX* | This study |
| yPN767 | *S. cerevisiae W303 can1-100 his3-11,15 ura3^0 BUD4-S288C COX4-2XGGGGGS-NFAST COX5A-MTS-CFAST10-2XGGGGGS-COX5A pet191Δ::HphMX* | This study |

| | | |
|---|---|---|
| yPN706 | *S. pombe h+ ade6-M210 leu1 bfr1::hygr pmd1::natr* | SAK27 from Kawashima et al (2012) |
| yPN671 | *S. cerevisiae W303 MAT**a** can1-100 ura3-1 trp1-1 tpk1-as tpk2-as tpk3-as pRS403-CGGR-pCGGRO-mCitrine-tADH1 (pPN309) pdr5Δ::KanMX* | This study |

**Table S2** *Plasmids used in the study*

| Plasmid ID | Vector and inserts | Source |
|---|---|---|
| pPN143 | *pRS403 with pRS403-pTDH3-yoPercevalHR-tADH1* | This study.<br><br>pTDH3-yoPercevalHR-tADH1 obtained from BYP9646 (NBRP) submitted by Nguyen et al. (2019). |
| pPN188 | *pRS403 with pCGGRO-mCitrine-tADH1* | This study.<br><br>pCggRO-mCitrine-tADH1 was amplified from Addgene ID 124582, submitted by Monteiro et al. (2019). |
| pPN309 | *pRS403 with pCGGRO-mCitrine-tADH1 and CGGR-R250A* | This study.<br><br>Constructed from pPN188 by inserting pTEFmut7-CggR (R250A)-tADH1 from Addgene ID 124585 submitted by Monteiro et al. (2019). |
| pPN382 | *pRS403 with pTEF1-Super ecliptic pHluorin-mRuby2-tADH1* | This study.<br><br>pTEF1-SEP-mRuby2-tADH1 was amplified from pKL08 obtained from Addgene ID 104432 submitted by Dodd et al. (2017) |
| pPN383 | *pMYT095 with Cox4 and Cox5a gRNA* | This study.<br><br>CRISPR plasmid for simultaneously tagging Cox4 and Cox5a with FAST fragments. |

**Movies**

**Movie S1** Timelapse movie of cells expressing PercevalHR and Superecliptic pHluorin which report ATP:ADP and cytosolic pH respectively. During the time course, the cells were moved from 2% glucose to 2% galactose followed by transfer back to 2% glucose. Arrestors and recoverers were assigned based on their growth recovery on galactose.

**Movie S2** Timelapse movie of cells expressing FBP reporter driven mCitrine and stained simultaneously with mitochondrial membrane potential dye TMRM (red). During the time course, the cells were moved from 2% glucose to 2% galactose.

**Movie S3** Timelapse movie of cells with mtDNA encoded and translated superfolder GFP ($sfGFP^{mt}$), and an mNeptune directed to mitochondrial matrix. Cells were grown in 2% glucose and shifted to 2% galactose.
